# Brain cholesterol metabolites cause significant neurodegeneration in human iPSC-derived neurons

**DOI:** 10.1101/2025.06.10.656594

**Authors:** Yuqing Feng, Bismoy Mazumder, Tasuku Konno, Ernestine Hui, Marius Brockhoff, Valentina Davi, Meng Lu, Edward Ward, Amberley Stephens, Wenyue Dai, Ana Fernandez-Villegas, Giuliana Fusco, Mohsen Ali Asgari, Edward Avezov, Alfonso De Simone, Yuqin Wang, William J Griffiths, Clemens Kaminski, Gabriele Kaminski Schierle

## Abstract

Disrupted cholesterol metabolism is increasingly recognised as a contributing factor in neurodegeneration; however, the specific effects of key brain-derived cholesterol metabolites, 24S-hydroxycholesterol (24S-HC) and 27-hydroxycholesterol (27-HC), remain poorly understood. Using human iPSC-derived i^3^ cortical neurons, we demonstrate that both 24S-HC and 27-HC significantly impair neuronal calcium signalling by elevating resting calcium levels, reducing spike amplitude, and disrupting network synchrony. These functional deficits are accompanied by widespread organelle dysfunction. Both oxysterols induce mitochondrial fragmentation, decrease spare respiratory capacity, and impair lysosomal degradation. Notably, 27-HC uniquely triggers lysosomal swelling and membrane permeabilisation. Additional signs of cellular stress, including axonal swellings and elevated endoplasmic reticulum calcium levels, were also observed. Furthermore, both 24S-HC and 27-HC were found to directly interact with alpha-synuclein (aSyn), promoting its accumulation in cellular models. In contrast, cholesterol itself had minimal impact, highlighting the distinct toxicity of its hydroxylated metabolites. Together, these findings reveal a mechanistic link between oxysterol accumulation and neuronal dysfunction, supporting the hypothesis that elevated levels of 24S-HC and 27-HC, commonly observed in Parkinson’s and Alzheimer’s disease, may actively drive neurodegenerative processes. Targeting oxysterol metabolism may therefore represent a promising therapeutic avenue for intervention in neurodegenerative disorders.

## Introduction

The brain contains the highest concentration of sterols in the body^1^, primarily in the form of cholesterol, along with its precursors 7-dehydrocholesterol and desmosterol^2^, and its metabolites 24S-hydroxycholesterol (24S-HC), and to a lesser extent, 27-hydroxycholesterol (27-HC, formally (25R)26-hydroxycholesterol)^3^. Cholesterol is mainly synthesised in astrocytes and, due to the blood–brain barrier (BBB), cannot cross into or out of the brain^4^. To maintain cholesterol homeostasis, excess cholesterol is transported to neurons and converted by the enzyme CYP46A1 into 24S-HC^5^, which, owing to its additional hydroxyl group, can exit the brain via circulation^1^. In contrast, 27-HC is predominantly produced in the liver by CYP27A1 and can enter the brain by crossing the BBB^6^. Under normal conditions, both 24S-HC and 27-HC help regulate cholesterol transport and maintain brain cholesterol homeostasis^7,8^.

Disruptions in cholesterol metabolism, particularly involving 24S-HC and 27-HC, have been implicated in neurodegenerative diseases^9,10^. Several factors may contribute to their elevated levels in the brain. As the brain ages, cholesterol accumulates, particularly due to processes like myelin breakdown, which in turn can elevate the production of 24S-HC^11^. At the same time, systemic metabolic changes, such as those induced by high-cholesterol diets, lead to increased hepatic synthesis of 27-HC, raising its levels in circulation and promoting its entry into the brain^12,13^. This influx is further exacerbated by age-associated breakdown of the BBB^14^, which becomes more permeable over time, allowing greater passage of peripheral 27-HC into the brain parenchyma^15,16^. Clinically, such dysregulation is reflected in elevated concentrations of 24S-HC in the cerebrospinal fluid (CSF) of individuals with Parkinson’s disease (PD)^15^ and Alzheimer’s disease (AD)^17,18^ and, alongside increased levels of 27-HC and its metabolites, in the CSF of PD patients^15,19^. These metabolites are not merely biomarkers: both have been implicated in neurotoxic processes, including alpha-synuclein (aSyn) aggregation in PD^20,21^. Moreover, 27-HC has been shown to contribute to amyloid-beta (Aβ) accumulation and tau phosphorylation, key hallmarks of AD pathology^22,23^.

Neuronal calcium signalling plays a central role in regulating synaptic plasticity and neurotransmission^24–26^. Intracellular calcium levels are tightly controlled through membrane channels and exchanges with internal stores, such as the endoplasmic reticulum (ER), mitochondria but also lysosomes^27,28^, which are frequently impaired in neurodegeneration^29–31^. Thus, understanding how cholesterol metabolites affect calcium dynamics and organelle health could uncover new mechanisms underlying neurodegenerative disease.

In this study, we utilised i^3^ cortical neurons derived from human induced pluripotent stem cells (iPSCs)^32^ to investigate the cellular and organelle-level effects of cholesterol, 24S-HC, and 27-HC. Using live-cell calcium imaging, structured illumination microscopy (SIM), and complementary biochemical assays, we show that 24S-HC and 27-HC impair calcium signalling and disrupt mitochondrial, lysosomal, and ER function. Given that cholesterol is known to directly interact with aSyn^33^, we further investigated whether these oxysterols influence aSyn dynamics. Our results demonstrate that both 24S-HC and 27-HC bind to aSyn with significantly higher affinity than cholesterol and promote its accumulation in cellular models. These findings establish a mechanistic link between cholesterol metabolites and neurodegenerative processes, particularly in the context of PD. They also highlight the potential of targeting elevated 24S-HC and 27-HC levels as a novel therapeutic strategy for mitigating aSyn pathology and disease progression.

## Methods

**Table 1.** Key reagents.

| Reagent or Resource | Source | Identifier |
| --- | --- | --- |
| **i<sup>3</sup> neuronal cell culture** |  |  |
| Stem Flex Medium | Gibco | A3349401 |
| Greiner CELLSTAR 6 well cell culture plates | Greiner Bio-One | 657160 |
| Vitronectin (VTN-N) Recombinant Human Protein, Truncated | Gibco | A14700 |
| StemPro™ Accutase™ Cell Dissociation Reagent | Gibco | A1110501 |
| Dulbecco's Modified Eagle Medium:Nutrient Mixture F-12 (DMEM / F-12), HEPES | Gibco | 11330032 |
| N-2 Supplement (100X), 5mL | Gibco | 17502-048 |
| Non-Essential Amino Acids Solution (100X) | Gibco | 11140050 |
| Antibiotic-Antimycotic (100X) | Gibco | 15240062 |
| 2-Mercaptoethanol (50 mM) | Gibco | 31350010 |
| Doxycycline hyclate | Sigma | D9891-1G |
| Geltrex™ LDEV-Free, hESC-Qualified, Reduced Growth Factor Basement Membrane Matrix | Gibco | A1413302 |
| Neurobasal medium 1X minus phenol red | Gibco | 12348017 |
| B-27™ Supplement (50X), serum free | Gibco | 17504044 |
| GlutaMAX™ Supplement | Gibco | 35050038 |
| Recombinant Human NT-3 (10ug) | Peprotech | 450-03-10UG |
| Human BDNF Recombinant Protein, 10 ug | Peprotech | 450-02 |
| ROCK Inhibitor (Y-27632) | BD Biosciences | 562822 |
| Nunc LabTek II Chambered Coverglass, 8 chambers (pack of 16) | Thermo Scientific | 155409PK |
| Corning® 96-well Flat Clear Bottom Black Polystyrene TC-treated Microplates, Individually Wrapped, with Lid, Sterile | Corning | 3603 |
| Poly-L-lysine solution, 0.01% | Sigma | P4707 |
| Incucyte® Neuroburst Orange Lentivirus (Synapsin) | Sartorius | 4736 |
| **COS-7 and HEK cell culture** |  |  |
| Dulbecco's Modified Eagle's Medium (DMEM) - high glucose | Sigma-Aldrich | D6546-500ML |
| Fetal Bovine Serum (FBS) | Gibco | A5256801 |
| **Plasmids and transfection reagents** |  |  |
| pLV_CMV_mEmerald-KDEL | Made by Dr Edward Avezov's group | - |
| ER-GCaMP-150 | Addgene | Plasmid #86918 |
| pMDLg_pRRE | Addgene | Plasmid #12251 |
| pRSV-Rev | Addgene | Plasmid #12253 |
| pMD2.G | Addgene | Plasmid #12259 |
| Lipofectamine 2000 Transfection Reagent | Invitrogen | 11668027 |
| Polyethyleneimine (PEI), linear, M.W. 25,000 | Thermo Scientific Chemicals | 043896.01 |
| PEG 8000 Powder, Molecular Biology Grade | Promega | V3011 |
| NaCl | Sigma-Aldrich | S9888 |

| **Cholesterols and Drugs** |  |  |
| --- | --- | --- |
| Cholesterol (ovine) | Avanti Polar Lipid | 700000P |
| 24(S)-Hydroxycholesterol | Sigma-Aldrich | SML1648 |
| 27-hydroxycholesterol | Sigma-Aldrich | SML2042 |
| Dimethyl sulfoxide (DMSO) | Sigma-Aldrich | D8418-100ML |
| U18666A | Sigma-Aldrich | 662015 |
| Bafilomycin A1, (V)-ATPase inhibitor-10 ug | Abcam | ab120497 |
| MG-132 | calbiochem | 474791 |
| Carbonyl cyanide-p-trifluoromethoxyphenylhydrazone (FCCP) | Abcam | ab120081 |
| **Dyes** |  |  |
| MitoTracker Deep Red | Invitrogen | M22426 |
| MitoTracker Orange | Invitrogen | M7510 |
| SiR-lysosome kit | Spirochrome | SC012 |
| DQ-Red BSA | Invitrogen | D12051 |
| **Seahorse assay** |  |  |
| Seahorse XF Cell Mito Stress Test Kit | Agilent | 103015-100 |
| Seahorse FluxPaks | Agilent | 103793-100 |

### i^3^ neuronal cell culture

i^3^ neurons were differentiated from human iPSCs as previously described^32^. Briefly, human iPSCs were cultured in StemFlex medium in 6-well cell culture plates with Vitronectin coating and split with Accutase while reaching 80 % confluence. After two or three passages, human iPSCs were differentiated in DMEM/F12, HEPES supplemented with 1X N-2 supplement, 1X non-essential amino acids, 1X antibiotic-antimycotic, 50 μM 2-mercaptoethanol and 1 μg/mL doxycycline hyclate in 6-well cell culture plates with Geltrex coating. After daily media change for 3 days, cells were split and transferred to Neurobasal medium supplemented with 1X B-27 supplement, 2 mM GlutaMAX supplement, 1X antibiotic-antimycotic, 50 μM 2-mercaptoethanol, 1 μg/mL doxycycline hyclate, 10 ng/mL NT-3, and 10 ng/mL BDNF. Full media was changed for four days followed by half media change every other day until day 21 in vitro (DIV). From 21 DIV onwards, full media was changed every other day. 10 μM ROCK inhibitor was added every time when plating cells. For different imaging purposes, 120,000 cells/well were plated in µ-slide 8 well high glass bottom plates (Nunc Lab-Tek II Chambered Coverglass), or 40,000-45,000 cells/well were plated in 96-well plates, all coated with 0.01% poly-l-lysine (PLL).

### COS-7 cell culture

COS-7 cells purchased from the American Type Culture Collection (ATCC) were cultured at 37°C with a 5% CO_2_ atmosphere. The cell culture media is composed of 90% Dulbecco′s Modified Eagle′s Medium-high glucose (DMEM), 10% fetal bovine serum (FBS), 2 mM GlutaMAX supplement, and 1X antibiotic-antimycotic. COS-7 cells were passaged when they reached 80-90% confluence.

### HEK cell culture and lentivirus production

Human embryonic kidney 293 (HEK293) cells purchased from ATCC were cultured at 37°C with a 5% CO_2_ atmosphere with the same media as COS-7 cells but without antibiotics. When the confluence reached 80-90%, HEK293 cells were transfected with mEmerald-KDEL plasmid, or ER-GCaMP6-150 using 3rd generation lentiviruses: pMDLg_pRRE, pRSV-Rev, pMD2.G. The transfection was performed using Lipofectamine 2000 transfection reagent or polyethyleneimine (PEI) with a plasmid ratio of 2:1:1:4 (pMDLg_pRRE: pRSV-Rev: pMD2.G: transfer plasmid). After 72 h of incubation, cell media was harvested with 5 minutes centrifugation at 500 x g and filtered with a 0.45 μm filter. To concentrate, lentiviruses were incubated with 4 x lentivirus concentrator solution (40% w/v PEG-8000 and 1.2 M NaCl in PBS, pH 7), shaking at 60 rpm overnight at 4 ℃, followed by spinning down at 1600 x g for 1 h. After removing the supernatant, the pellet was resuspended in PBS and aliquoted to working volume.

### Cholesterol, 24S-HC and 27-HC preparation

Cholesterol (ovine), 24S-HC and 27-HC were dissolved in dimethyl sulfoxide (DMSO) or ethanol to a stock concentration of 5 mM and aliquoted to amber glass vials to store at −20 ℃.

### Cholesterol, 24S-HC, 27-HC and drug treatment

Unless specified, i^3^ neurons (26-27 DIV) were treated for 3 days with 10 μM cholesterol, 10 μM 24S-HC or 10 μM 27-HC with replenishment when changing the media. For lysosomal morphology treatment, 10 μM U18666A was added for 3 days as a positive control.

### Organelle staining

#### Mitochondria

On the day of imaging, mitochondria were stained with 50 nM MitoTracker Deep Red or MitoTracker Orange CMTMRos for 30 minutes, followed by washing the cells once with culture media.

#### Lysosome

The day before imaging, i^3^ neurons were stained with 1.5 μM SiR-lysosome and 10 μM verapamil hydrochloride solution and incubated overnight. On the day of imaging, cells were washed twice with the culture media. 10 μM verapamil hydrochloride solution was added again with fresh culture media.

#### ER

mEmerald-KDEL lentivirus and ER-GCaMP6 lentivirus were added to i^3^ neurons at 14-15 DIV to monitor the ER morphology and ER calcium level, respectively.

### Widefield microscopy

Widefield imaging was performed with an automated custom-built widefield microscope. 4-wavelength high-power LED source (LED4D067, Thorlabs), stage (Prior), frame (IX83, Olympus), Z drift compensator (IX3-ZDC2, Olympus) and an sCMOS camera (Zyla sCMOS, Andor) were controlled by Micro-Manager. Filter cubes for ER-GCaMP6 (filter set 49002-ET-EGFP, Chroma), calcium imaging or DQ-Red BSA (filter set 49008-ET-mCherry, Texas Red, Chroma) were used. For calcium imaging, ER-GCaMP6 or DQ-BSA assay, images were captured with a 10×/0.25 air (Plan N, Olympus), a 20×/0.45 NA air (LUCPlanFL N, Olympus), or a 60×/1.42 NA oil (PlanApo U, Olympus) objective lens, respectively. Live-cell images were captured while keeping the cells at 37℃ with 0.5% CO2 atmosphere in a microscope stage top micro-incubator (OKO Lab).

### Structured illumination microscopy

The custom-made three-color SIM system was built on an Olympus IX71 microscope stage. Fluorophores were excited by lasers at the wavelengths of 488 nm (iBEAM-SMART-488, Toptica), 561 nm (OBIS 561, Coherent), or 640 nm (MLD 640, Cobolt). Structured illumination pattern was achieved by expanding the laser beam to fill the display of a ferroelectric binary Spatial Light Modulator (SLM) (SXGA-3DM, Forth Dimension Displays). A Pockels cell (M350-80-01, Conoptics) was used to control the polarisation of light. After focusing the structured illumination pattern onto the sample and collecting the emission light with a 60×/1.2 numerical aperture (NA) water immersion lens (UPLSAPO 60XW, Olympus), images were captured by a sCMOS camera (C11440, Hamamatsu). HCImage software (Hamamatsu) and a custom-made LabView program worked synchronously to acquire raw images. Live-cell SIM images were captured while keeping the cells on the OKO Lab stage. SIM images were reconstructed with LAG SIM, a custom-made ImageJ plugin after calibration of channel displacement with TetraSpeck beads (Life Technologies).

### Calcium imaging

Incucyte Neuroburst Orange lentivirus (Sartorius) was added to i^3^ neurons at about 14-15 DIV with 2 μL/20,000 cells. From about 26 DIV, calcium imaging was performed with widefield microscopy which is described earlier in this section. Neurons (26-27 DIV) were treated for 4 days with 10 μM cholesterol, 10 μM 24S-HC, 10 μM 27-HC, 3 nM bafilomycin A1, 140 nM MG-132 or 2 μM carbonyl cyanide-p-trifluoromethoxyphenylhydrazone (FCCP) after pretreatment imaging. The treatment was first added on Day 0 when the media was changed and replenished on Day 2 after imaging during the next media change to maintain consistent exposure. The same volume of DMSO or ethanol was added as vehicle control. Imaging was performed once daily over a period of five consecutive days. 1000 frames were captured with 3*3 binning, 100 ms exposure time and 0 intervals with an overall imaging time of 126 seconds.

### Seahorse Mito Stress Tests

10,000 cells per well of COS-7 cells were plated in the 96-well plate (XF96). 24 h after seeding, cells were treated with vehicle, 10 μM cholesterol, 10 μM 24S-HC or 10 μM 27-HC daily for two days. 1.5 μΜ oligomycin, 1 μΜ FCCP and 0.5 μΜ rotenone/antimycin A (Seahorse XF Cell Mito Stress Test Kit) were sequentially added to the cells during the oxygen consumption rates (OCR) measurement performed by a Seahorse XFe96 analyser (Agilent). Data were normalised to the cell number and analysed on Wave (Agilent). The spare respiratory capacity was calculated by subtracting the normalised OCR at time point 3 from time point 9.

### DQ-BSA assay

i^3^ neurons transduced with mEmerald-KDEL lentivirus were stained with 10 µg/mL DQ Red BSA for 6 h, followed by washing with PBS twice. In the meantime, 0.5 μM bafilomycin A1 was added for 6 h as a negative control. i^3^ neurons were imaged using widefield microscopy which is described earlier in this section while keeping the cells on the OKO Lab stage. Images were captured with an exposure time of 100 ms. To avoid photo bleaching, somas were focused with the ER channel followed by the capture of both channels. DQ-BSA assay data analysis was performed using imageJ. The ER channel was used to identify the soma by applying Otsu’s thresholding and using the “Analyse Particles” function in imageJ to get the region of interest (ROI). The DQ-BSA fluorescence intensity of the soma was measured by applying the saved ROI after subtracting the background noise.

### Expression and purification of ^15^N-labelled recombinant aSyn for NMR studies

Recombinant aSyn was expressed in *E. coli* using the plasmid pT7-7 as previously described^34^. In order to obtain N-terminal acetylation of aSyn, coexpression with a plasmid carrying the components of the NatB complex (Addgene) was used. After transforming into BL21-Gold (DE3) cells (Agilent Technologies), uniformly ^15^N labelled aSyn was obtained by growing the bacteria in isotope-enriched M9 minimal media containing 1 g/L of ^15^N ammonium chloride (Sigma-Aldrich) and 1 g/L of ^15^N IsoGro (Sigma). Cell growth was performed at 37 °C under constant shaking at 250 rpm and using 100 μg/mL ampicillin. Upon reaching an OD600 of 0.6, aSyn expression was induced with 1 mM isopropyl β-D-1-thiogalactopyranoside (IPTG) at 37 °C for 4 h, and subsequently cells were harvested by centrifugation at 6,200 x g (Beckman Coulter). Cell pellets were then resuspended in lysis buffer (10 mM Tris-HCl pH 8, 1 mM EDTA and EDTA-free complete protease inhibitor cocktail tablets (Roche) and lysed by sonication. The cell lysate was centrifuged at 22,000 x g for 30 minutes to remove cell debris. In order to precipitate heat-sensitive proteins, the supernatant was then heated for 20 minutes at 70 °C and centrifuged at 22,000 x g. Subsequently streptomycin sulfate was added to the supernatant to a final concentration of 10 mg/mL to stimulate DNA precipitation. The mixture was stirred for 15 minutes at 4 °C followed by centrifugation at 22,000 x g. Subsequently, ammonium sulfate was added to the supernatant to a concentration of 360 mg/mL in order to precipitate the protein, and the solution was then stirred for 30 minutes at 4 °C prior to further centrifugation at 22,000 x g. The resulting pellet was resuspended in 25 mM Tris-HCl, pH 7.7 and dialysed against the same buffer in order to remove salts. The dialysed solutions were then loaded onto an anion exchange column (26/10 Q sepharose high performance, GE Healthcare) and eluted with a 0–1 M NaCl step gradient, and then further purified by loading onto a size exclusion column (Hiload 26/60 Superdex 75 preparation grade, GE Healthcare). All the fractions containing the monomeric protein were pooled together and concentrated by using Vivaspin filter devices (Sartorius Stedim Biotech). The purity of the aliquots after each step was analysed by SDS-PAGE and the protein concentration was determined from the absorbance at 275 nm using an extinction coefficient of 5600 M^-1^ cm^-1^. Mass spectrometry was used to assess that the level of N-terminal acetylation was complete.

### NMR

NMR experiments were carried out at 10 °C on a Bruker spectrometer operating at a ^1^H frequency of 700 MHz equipped with triple resonance HCN cryo-probe, and data were collected using TopSpin 4.4.0 software (Bruker, AXS GmBH). The aSyn concentration was kept constant at 10 µM in all experiments. ^1^H-^15^N HSQC spectra were recorded using a data matrix consisting of 2048 (t2, 1H) × 220 (t1, 15N) complex points. Assignments of the resonances in ^1^H-^15^N HSQC spectra of aSyn are derived from our previous studies, and data were analysed using Sparky 3.1 software.

### Data analysis

#### Calcium imaging

The calcium imaging data were analysed using a custom Python script. Maximum intensity projection images were generated from time-series fluorescence images, followed by Otsu’s thresholding to create a binary mask. Individual somas were identified using rectangular bounding boxes applied to the binarized images, with adjustable size parameters to exclude clustered or improperly segmented regions. Raw calcium traces were extracted from each selected soma and normalised to ΔF/F_0_ calculated using the formula:

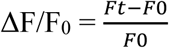

where F_t_ is the fluorescence intensity at each time point, and F_0_ represents the baseline fluorescence intensity, indicative of the resting calcium level. It is determined by averaging the fluorescence intensity over a designated time window in which no calcium spikes are observed, ensuring a stable baseline measurement. For each trace, relative calcium spike amplitudes (peak ΔF/F_0_) were computed by normalising to the baseline fluorescence intensity within the trace:

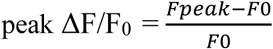

where F_0_ is the baseline fluorescence intensity, and F_peak_ is the peak fluorescence intensity. The number of detected peaks was counted and normalised by the total recording time to determine the spike rate (spikes per second). To quantify calcium spike synchronicity, we computed the Spike Time Tiling Coefficient (STTC), a metric designed to assess the temporal alignment of spike events between calcium traces^35^. For each pair of detected calcium traces, the STTC was calculated using a time window (±Δt) of 0.5 seconds. This metric compares the total proportion of recording time that falls within the time window around spikes in one trace to the proportion of spikes in another trace that also lie within that time window. The STTC is computed as follows:

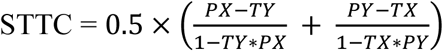

where PX and PY are the proportions of spikes in traces X and Y that fall within ±Δt of a spike in the other trace, and TX and TY represent the fraction of total recording time that lies within ±Δt of any spike in traces X and Y, respectively.

### Mitochondrial morphology

Mitochondrial morphology was analysed using ImageJ. To do the segmentation of mitochondria, Otsu’s thresholding was applied to each image, followed by the “Analyse Particles” function in imageJ with a size filter excluding objects smaller than 0.1 μm². The circularity of each mitochondrion was calculated automatically by ImageJ using the formula: 4π(area/perimeter^2^). The value of circularity is ranged from 0 to 1, where 1 indicates a perfect circle.

### Lysosomal morphology

Lysosomal morphology was analysed using ImageJ. To do the segmentation of lysosomes, Otsu’s thresholding was applied to each image, followed by the “Analyse Particles” function in imageJ with a size filter excluding objects smaller than 0.1 μm². The area of each lysosome was measured automatically by ImageJ. The number of neurons that have lysosomal dye leakage was counted manually.

### ER calcium

After subtracting the background noise, Otsu’s thresholding was applied to each image, followed by the “Analyse Particles” function in imageJ with a size filter to identify the somas. The ER-GCaMP6 fluorescence intensity was measured for each ROI.

### Statistical analysis

All statistical analysis was done using GraphPad Prism 8 or 10.

## Results

### 24S-HC and 27-HC impair calcium signalling in i^3^ neurons

We first investigated how 24S-HC and 27-HC affect neuronal function at the cellular level using human iPSC-derived i^3^ neurons. To determine the optimal treatment concentration, we employed calcium imaging, as calcium signalling serves as a key indicator of neuronal function and activity. For this, i³ neurons were transduced with the Incucyte Neuroburst Orange lentivirus, a cytosolic calcium indicator, and imaged using widefield microscopy at approximately day 26 in vitro (DIV), a time point characterised by spontaneous, synchronised calcium activity^36^, indicative of functional network maturation (Figure S1). Following baseline imaging, neurons were treated with 2.5, 5, or 10 µM of cholesterol, 24S-HC, or 27-HC for four days, and imaged daily over a five-day period. For each time-series of fluorescence images, individual somas were identified via thresholding to plot the cytosolic calcium traces, while clustered neurons were filtered out (Figure 1a). Our results show that both 24S-HC and 27-HC impaired calcium signalling in a dose-dependent manner, whereas cholesterol did not exhibit a clear trend (Figures S2–S4). Based on these findings, we selected 10 µM for further experiments, as it produced a clear biological response without excessive cytotoxicity.

**Figure 1.**
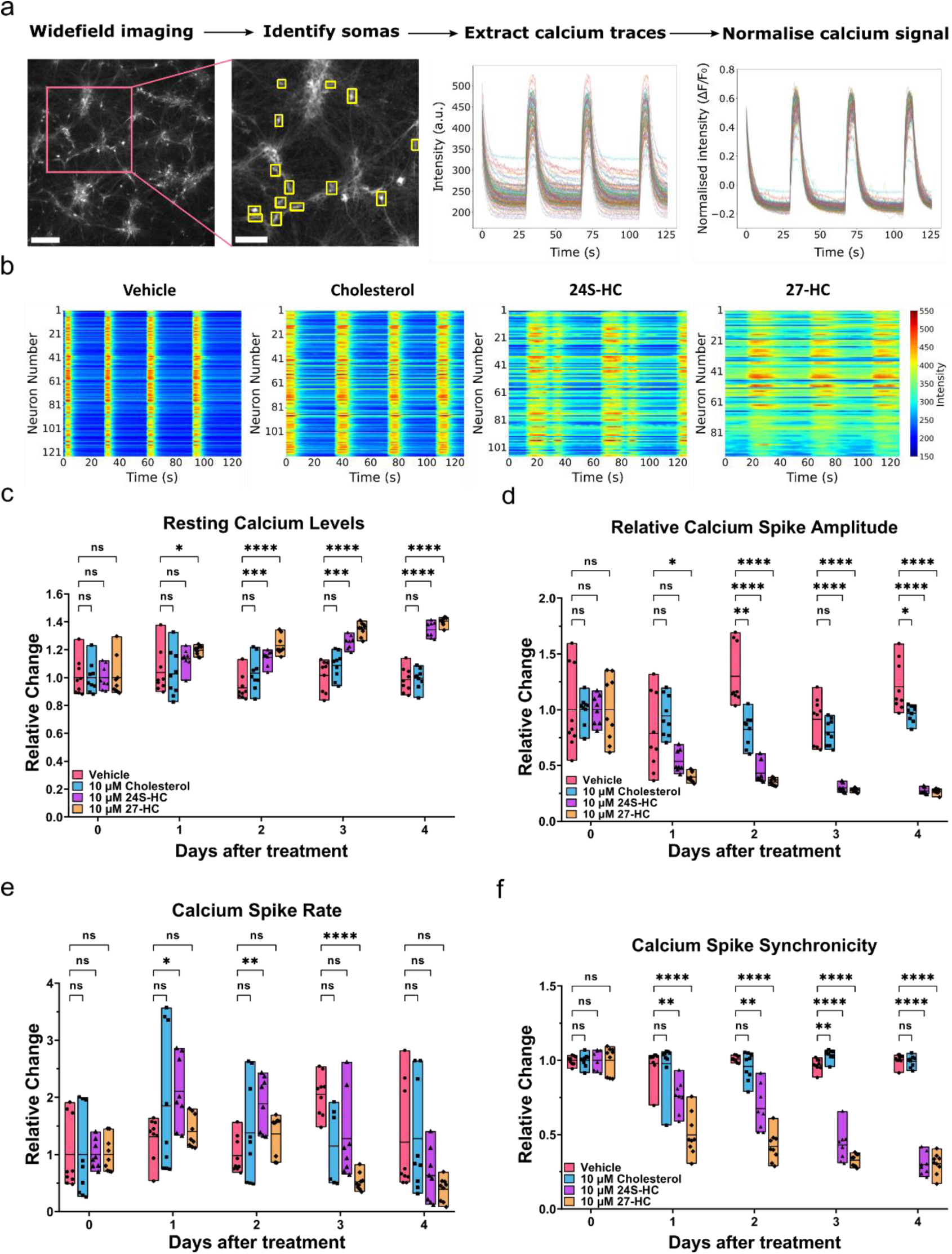
24S-HC and 27-HC impair calcium signalling in i^3^ neurons. **(a)** A brief overview of the calcium imaging methodology and subsequent data analysis. i^3^ neurons transduced with the Incucyte Neuroburst Orange lentivirus were imaged using widefield microscopy, scale bar = 200 μm. Individual somas were identified (yellow rectangles) after thresholding, scale bar = 100 μm. The fluorescence intensity fluctuation of each soma over time was extracted, then normalised to ΔF/F_0_. **(b)** Calcium imaging heatmaps showing activity of i^3^ neurons treated with 10 µM cholesterol, 24S-HC or 27-HC for 1 day. **(c) - (f)** The relative change in resting calcium levels (F_0_), relative calcium spike amplitude (peak ΔF/F_0_), calcium spike rate and calcium spike synchronicity of i^3^ neurons treated with 10 µM cholesterol, 24S-HC or 27-HC over 4 days. Data were normalised to Day 0 (pretreatment) and presented as floating bar plots (min to max), with a line indicating the mean. One field of view was imaged per well, and each data point represents a single well. **(c)**, **(d)** and **(f)** were analysed with two-way ANOVA using Dunnett’s multiple comparisons tests, while **(e)** was analysed with mixed-effects analysis using Dunnett’s multiple comparisons tests. ns is not significant, * is p<0.05 and **** is p<0.0001. Data from N = 3 independent experiments were quantified.

After one day of treatment, both 10 µM 24S-HC and 27-HC altered the calcium firing patterns in i³ neurons compared to vehicle controls (Figure 1b). These changes were characterised by a broadening of calcium spikes and elevated basal intracellular calcium levels. To further characterise these effects, we analysed calcium dynamics over a five-day period, including resting calcium levels, relative calcium spike amplitude, spike rate, and synchronicity. 27-HC significantly increased resting calcium levels (F_0_) after just one day of treatment, whereas the effect of 24S-HC emerged after two days and continued to increase progressively, indicating a sustained elevation in cytosolic calcium level (Figure 1c). In parallel, 27-HC significantly reduced the relative calcium spike amplitude (peak ΔF/F_0_) after one day, with 24S-HC producing a comparable reduction after two days (Figure 1d). These reductions worsened over time, reflecting a progressive dampening of intracellular calcium transients. Interestingly, 24S-HC transiently increased spike rate during the first two days, while 27-HC led to a significant decrease in spike rate from day three onwards (Figure 1e). Although the temporal patterns differed, both oxysterols disrupted the normal frequency and regulation of calcium activity. Notably, both compounds significantly reduced calcium spike synchronicity from day one, with this effect becoming increasingly pronounced over time (Figure 1f), suggesting impaired coordinated activity across neuronal networks. In contrast, cholesterol treatment did not produce consistent or significant changes in any of the measured calcium dynamics parameters (Figure 1c–f), likely due to its limited membrane permeability compared to 24S-HC and 27-HC, both of which contain an additional hydroxyl group facilitating cellular uptake^1^. Together, these findings demonstrate that treatment with 10 µM 24S-HC and 27-HC leads to significant, progressive disruption of calcium homeostasis and network activity in i³ neurons, highlighting their potential neurotoxic effects.

Several organelles are implicated in calcium homeostasis. To investigate which organelle/pathway is linked to the above observed calcium signalling effect, we treated the neurons with pharmacological inhibitors: 3 nM bafilomycin A1, 140 nM MG-132, and 2 µM carbonyl cyanide-p-trifluoromethoxyphenylhydrazone (FCCP), effective concentrations without excessive cytotoxicity. Bafilomycin A1 is an autophagy inhibitor that disrupts V-ATPase-dependent acidification and Ca-P60A/SERCA-mediated autophagosome-lysosome fusion^37^. MG-132 inhibits the ubiquitin-proteasome system^38^. FCCP is an uncoupler of oxidative phosphorylation in mitochondria, disrupting the proton gradient across the inner mitochondrial membrane, thereby reducing the driving force for ATP synthesis^39^. Our results show that treatment with 3 nM bafilomycin A1 for two days significantly increased resting calcium levels, whereas 2 µM FCCP led to a reduction (Figure S5a). In contrast, 140 nM MG-132 did not produce a significant effect on resting calcium levels. All three treatments affected the relative calcium spike amplitude: bafilomycin A1 and MG-132 induced reductions after three and two days, respectively, while FCCP caused a decrease after either one or three days of exposure (Figure S5b). Additionally, calcium spike rate was significantly reduced following two days of treatment with each inhibitor, with progressive worsening over subsequent days, except in the case of FCCP (Figure S5c). A loss of calcium spike synchronicity was also observed after three or four days of treatment with MG-132 and bafilomycin A1, while FCCP did not significantly affect synchronicity (Figure S5d). In summary, these findings indicate that disruption of lysosomal function, proteostasis, or mitochondrial activity can each contribute to impaired calcium signalling in i^3^ neurons.

### 24S-HC and 27-HC cause mitochondrial dysfunction

Previous experiments showed that 10 µM 24S-HC and 27-HC induced significant calcium signalling impairments within 1 day, with effects worsening over time. To capture a representative stage of this phenotype while ensuring experimental consistency, we selected a 3-day treatment duration for subsequent experiments. Given the similarity in reduced relative calcium spike amplitude observed with 24S/27-HC and FCCP, we investigated whether 24S-HC and 27-HC contribute to mitochondrial dysfunction. As an initial indicator of potential alterations, we assessed mitochondrial morphology in i^3^ neurons treated with 10 µM cholesterol, 24S-HC, or 27-HC for 3 days, using SIM. Compared to vehicle control, 24S-HC or 27-HC treatment induced mitochondrial fragmentation in i^3^ neurons, while cholesterol had no significant impact (Figure 2a). Quantitative analysis confirmed a significant increase in mitochondrial circularity with 24S-HC and 27-HC but not cholesterol (Figure 2b). The same effect was also observed in COS-7 cells, the African green monkey kidney fibroblast-like cells, with 10 µM 24S-HC or 27-HC treatment for 1 day (Figure S6a and b), which we later use to run the Seahorse Mito Stress Test on.

**Figure 2.**
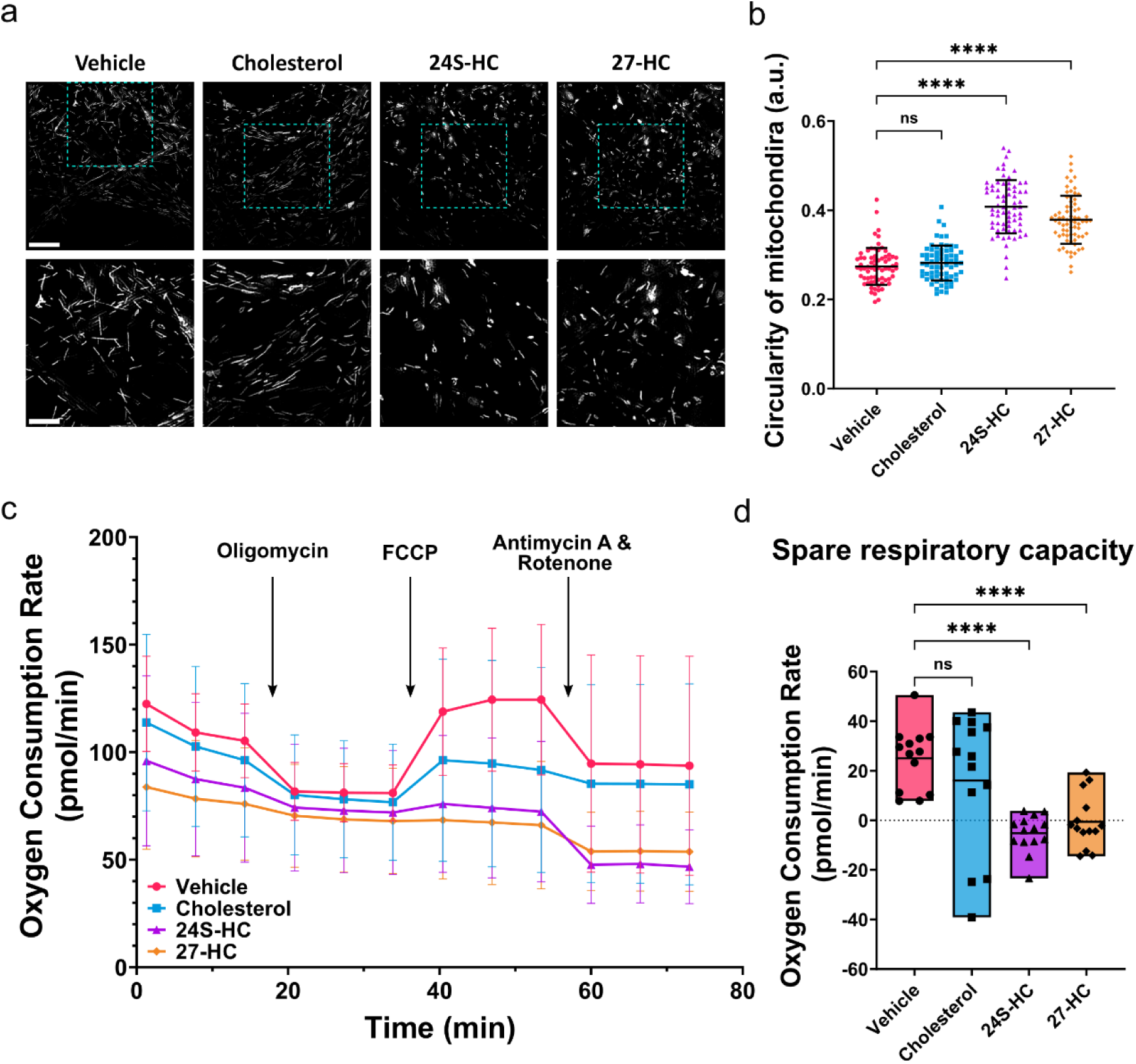
24S-HC and 27-HC cause mitochondrial dysfunction. **(a)** Top row shows representative SIM images of mitochondria in i^3^ neurons treated with 10 µM cholesterol, 24S-HC or 27-HC for 3 days; scale bar = 10 μm. Bottom row shows the zoomed-in area indicated by the cyan rectangle; scale bar = 5 µm. **(b)** The circularity of mitochondria is presented as means ± SD. Each data point represents an individual image. Kruskal-Wallis test with Dunn’s multiple comparisons, where ns is not significant and **** is p<0.0001. Data from N = 3 independent experiments were quantified. **(c)** Seahorse Mito Stress Tests show the oxygen consumption rate (OCR) level change of COS-7 cells treated with 10 µM cholesterol, 24S-HC or 27-HC daily for two days. Data were normalised to the value of the untreated sample in each repeat. **(d)** The spare respiratory capacity was presented as floating bar plots (min to max), with a line indicating the mean. Each data point represents an individual well. Brown-Forsythe ANOVA test with Dunnett’s T3 multiple comparisons tests, where ns is not significant and **** is p<0.0001. Data from N = 3 independent experiments were quantified.

Since structural changes often precede or accompany functional impairments, we assessed mitochondrial respiration using the Seahorse Mito Stress Tests. COS-7 cells were treated daily with 10 µM cholesterol, 24S-HC or 27-HC for two days, and the oxygen consumption rate (OCR) was measured upon the addition of oligomycin, FCCP and antimycin A and rotenone (Figure 2c). After FCCP addition, the spare respiratory capacity was significantly reduced in COS-7 cells treated with 24S-HC or 27-HC, but not with cholesterol, compared to vehicle control (Figure 2d), indicating impaired mitochondrial respiration. Collectively, these results show that 24S-HC and 27-HC induce mitochondrial dysfunction in i^3^ neurons and COS-7 cells.

### 24S-HC and 27-HC cause lysosomal dysfunction in i^3^ neurons

Since 24S-HC, 27-HC and bafilomycin A1 induced similar calcium signalling defects, we next investigated whether 24S-HC and 27-HC affect lysosomal function. Using SIM, we examined lysosomal morphology in i^3^ neurons treated with 10 µM cholesterol, 24S-HC, or 27-HC for 3 days. U18666A, a drug that induces cholesterol accumulation in lysosomes and mimics Niemann-Pick disease type C (NPC), was used as a positive control^40^. Mitochondria were co-stained to identify somas. We found that 27-HC caused an enlargement of somatic lysosomes in i^3^ neurons, while neither cholesterol nor 24S-HC had a similar effect (Figure 3a). Quantification results further proved this, showing an increased lysosome size upon treatment with 27-HC (Figure 3b). The same effect was also observed in COS-7 cells (Figure S6c and d). Notably, 27-HC not only enlarged lysosomes but also induced lysosomal membrane permeabilisation (LMP), as evidenced by leakage of the lysosomal labelling dye (Figure 3c, orange arrows). The fluorescent traces likely reflect the trajectory of lysosome movement, as the leakage occurred while lysosomes were actively trafficking within the cell. Quantification results show that 76% of i^3^ neurons treated with 27-HC exhibited dye leakage, a phenomenon not observed in any other treatment conditions (Figure 3d).

**Figure 3.**
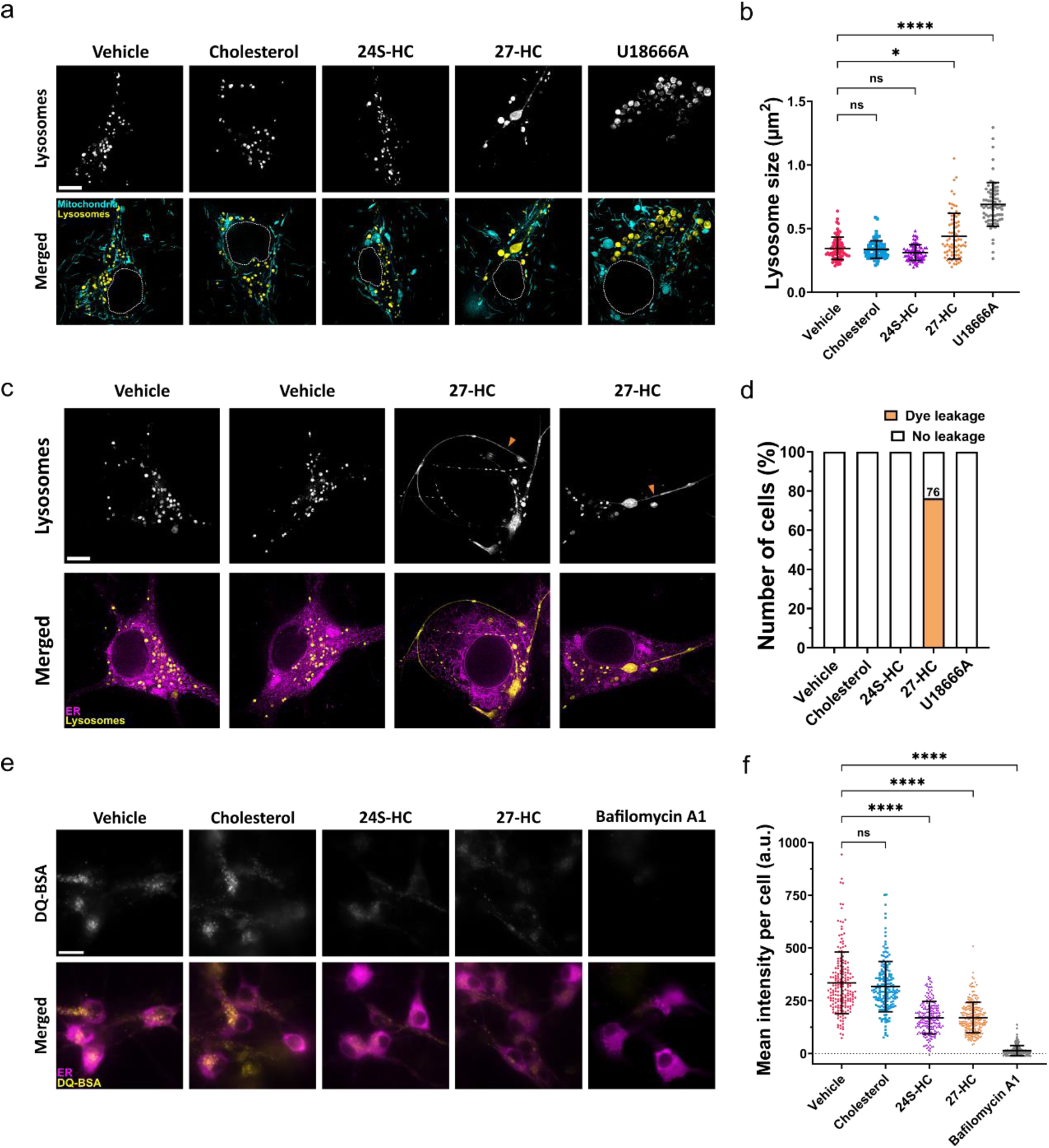
24S-HC and 27-HC cause lysosomal dysfunction in i^3^ neurons. **(a)** Top row shows representative SIM images of lysosomes in i^3^ neurons treated with 10 µM cholesterol, 24S-HC or 27-HC for 3 days. 10 µM U18666A was added for 3 days as a positive control. Bottom row: Mitochondria were co-stained to identify the soma, and the nucleus was manually outlined in white to indicate its location. Scale bar = 5 μm. **(b)** The size of lysosomes is presented as means ± S.D. Each data point represents an individual image. Kruskal-Wallis test with Dunn’s multiple comparisons, where ns is not significant, * is p<0.05 and **** is p<0.0001. Data from N = 3 independent experiments were quantified. **(c)** Top row shows representative SIM images of lysosomes in i^3^ neurons treated with 10 µM 27-HC, showing leakage of SiR-lysosome dye (orange arrows). Scale bar = 5 μm. Bottom row: the ER was labelled with an mEmerald-KDEL lentivirus to identify the soma. **(d)** The percentage of cells with dye leakage was plotted. Data from N = 3 independent experiments were quantified. **(e)** Representative widefield images of DQ-BSA fluorescence in i^3^ neurons treated with 10 µM cholesterol, 24S-HC or 27-HC for 3 days. The ER was labelled to identify the soma. 0.5 µM bafilomycin A1 was added for 6 h as a negative control. Scale bar = 20 μm. **(f)** The intensity of DQ-BSA in each soma is presented as means ± S.D. Each data point represents an individual image. Representative images in **(e)** are shown at higher magnification to enhance visibility and do not reflect the full field of view used for analysis. Kruskal-Wallis test with Dunn’s multiple comparisons, where ns is not significant and **** is p<0.0001. Data from N = 3 independent experiments were quantified.

To determine whether LMP leads to lysosomal dysfunction, we performed the DQ-BSA (dye quenched-bovine serum albumin) assay. In this assay, BODIPY-conjugated BSA remains self-quenched until lysosomal degradation of BSA releases the dye from its quenched state. i^3^ neurons were treated with 10 µM cholesterol, 24S-HC, or 27-HC for 3 days, with 0.5 µM bafilomycin A1 added for 6 h as a negative control. ER was labelled by transducing neurons with an mEmerald-KDEL lentivirus to identify the soma. Representative widefield images show markedly reduced fluorescence intensity in i^3^ neurons treated with 24S-HC and 27-HC, but not with cholesterol, compared to vehicle control (Figure 3e). Quantitative analysis confirmed a significant decrease in fluorescence intensity (Figure 3f), indicating impaired lysosomal degradation capacity. Together, these findings demonstrate that 24S-HC and 27-HC induce lysosomal dysfunction in i^3^ neurons.

### 24S-HC and 27-HC cause ER disturbances in i^3^ neurons

Since lysosomes regulate the tubular distribution of the ER^41^, the observed lysosomal impairment may contribute to ER dysfunction. To investigate this, i^3^ neurons were treated with 10 µM cholesterol, 24S-HC, or 27-HC for 3 days, and the ER, mitochondria, and lysosomes in axons near the soma were imaged using SIM. With 24S-HC and 27-HC treatment, ER structures appeared constricted and displaced in swellings of axonal regions, colocalised with enlarged mitochondria but not with lysosomes (Figure 4a, orange arrows). Quantification revealed a 4-to-5-fold increase in the number of swellings in axons with 24S-HC and 27-HC treatment compared to vehicle control and cholesterol-treated conditions (Figure 4b), suggesting that while axonal swelling may occur physiologically, it is markedly exacerbated under pathological conditions.

**Figure 4.**
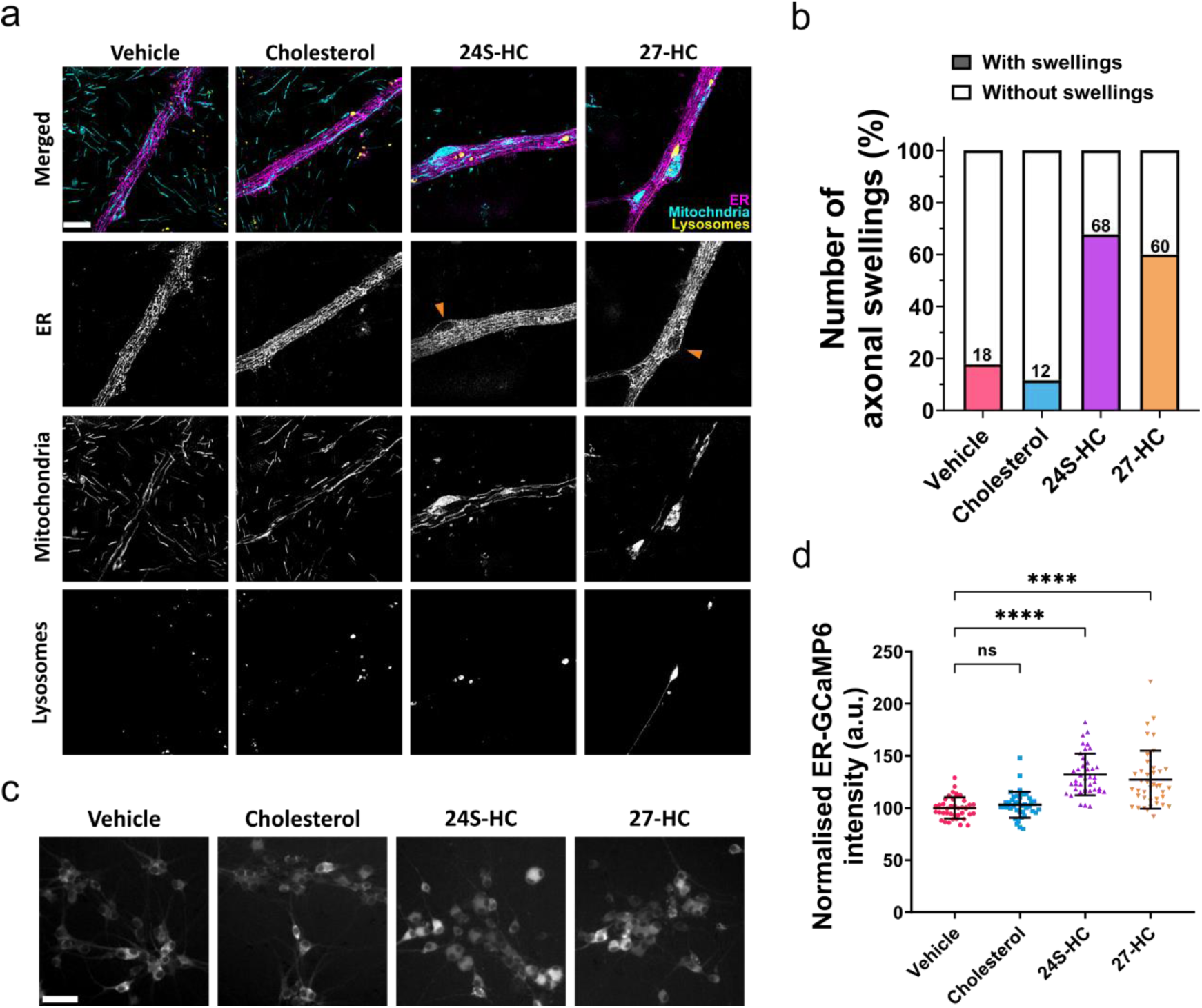
24S-HC and 27-HC cause ER disturbances in i^3^ neurons. **(a)** Representative SIM images of the ER, mitochondria and lysosomes in the axons of i^3^ neurons treated with 10 µM cholesterol, 24S-HC or 27-HC for 3 days. Scale bar = 4 μm. Axonal swellings were observed in 24S-HC and 27-HC treated conditions (orange arrows). **(b)** The percentage of axons with swelling was plotted. Data from N = 4 independent experiments were quantified. **(c)** Representative widefield images of ER-GCaMP6 fluorescence in i^3^ neurons treated with 10 µM cholesterol, 24S-HC or 27-HC for 3 days. Scale bar = 50 μm. **(d)** The normalised intensity of ER-GCaMP6 is presented as means ± S.D. Each data point represents an individual image. Representative images in **(c)** are shown at higher magnification to enhance visibility and do not reflect the full field of view used for analysis. Kruskal-Wallis test with Dunn’s multiple comparisons, where ns is not significant and **** is p<0.0001. Data from N = 3 independent experiments were quantified.

The ER is central to neuronal calcium homeostasis, regulating calcium storage and release, particularly during neuronal activity^42^. To assess ER calcium levels, i^3^ neurons were transduced with an ER-GCaMP6 lentivirus and imaged using widefield microscopy (Figure 4c). Quantification shows significantly higher ER-GCaMP6 fluorescence intensity in the soma of i^3^ neurons treated with 24S-HC and 27-HC, but not with cholesterol, compared to vehicle control (Figure 4d), indicating elevated ER calcium levels. Together, these findings demonstrate that 24S-HC and 27-HC induce ER disturbances in i^3^ neurons.

### 24S-HC and 27-HC bind directly to the N-terminus of aSyn with greater affinity than cholesterol

To further explore the connection between oxysterols and neurodegeneration, we investigated their role in the context of PD, a well-characterised neurodegenerative disorder marked by aSyn pathology. Previous studies have shown that both 24S-HC and 27-HC promote aSyn aggregation, as demonstrated by thioflavin T (ThT) fluorescence assays and cellular models^20,21^. Furthermore, cholesterol has previously been shown to directly interact with aSyn^33^. To further investigate whether the cholesterol metabolites 24S-HC and 27-HC also engage in direct interactions with aSyn, we performed nuclear magnetic resonance (NMR) spectroscopy. Our results reveal that both 24S-HC and 27-HC bind directly to the N-terminal region of aSyn, exhibiting significantly greater affinity than cholesterol (Figure 5).

**Figure 5.**
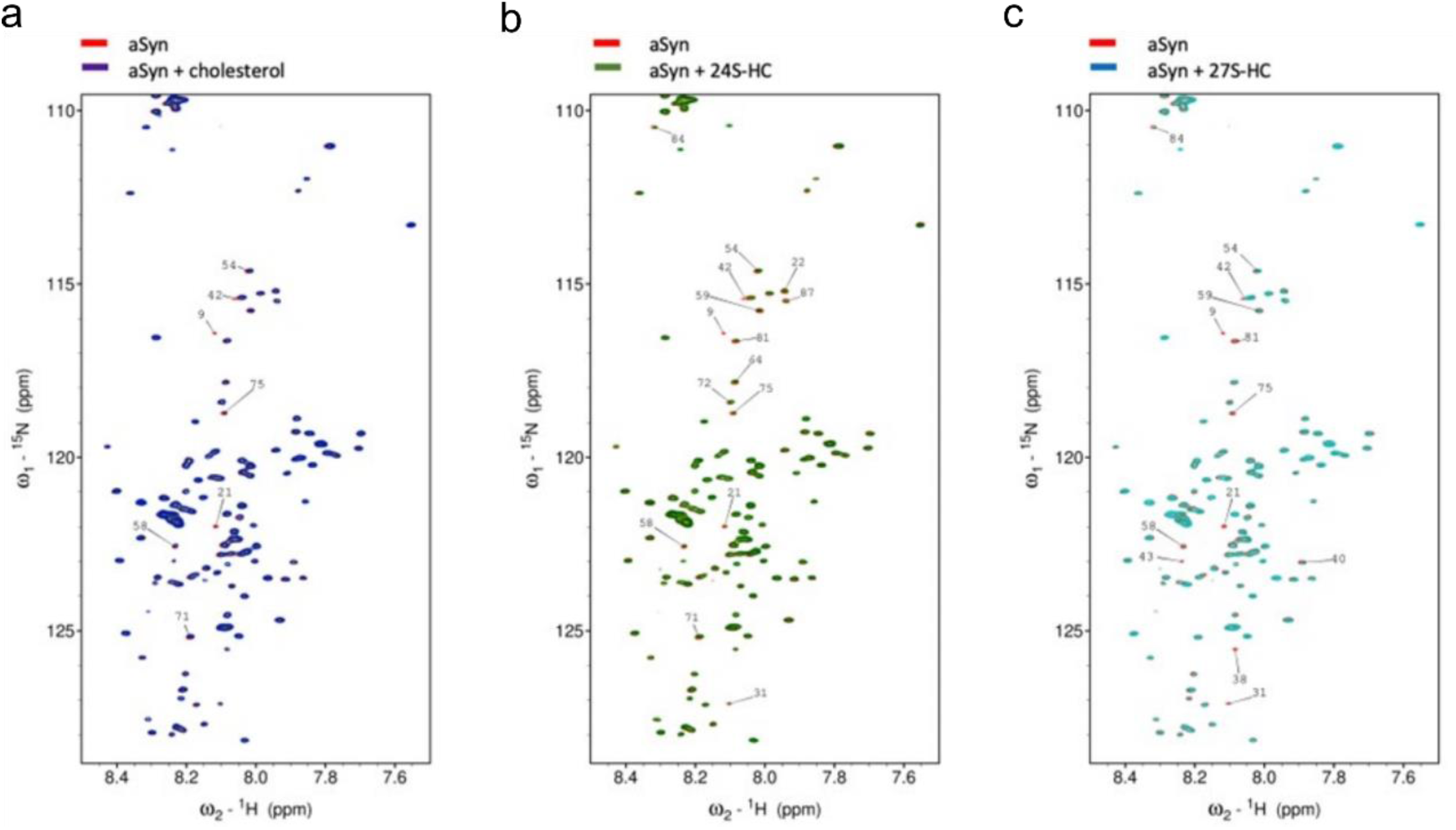
^1^H-^15^N HSQC NMR spectra detected chemical shift perturbations (CSP) of the resonances of monomeric aSyn (10 µM) in the presence of three molar equivalents of cholesterol, 24S-HC or 27-HC (30 µM). **(a)** A few of CSP in the ^1^H-^15^N resonances of aSyn amide backbone were found in the presence of cholesterol. These are primarily located in the membrane binding region at the N-terminal 90 residues (residues 9, 21, 42, 54, 58, 71, 75). Stronger effects were found in the case of 24S-HC **(b)** and 27-HC **(c)**, with CSP extending to additional residues (e.g., 22, 31, 38, 40, 43, 64, 81, 84, 87).

To examine the functional consequences of this interaction in a cellular context, we utilised SH-SY5Y neuroblastoma cells overexpressing aSyn-YFP. Cells were treated with 10 µM cholesterol, 24S-HC, or 27-HC for 2 days to assess their impact on aSyn levels. Our results show significantly increased YFP fluorescence intensity in cells treated with 24S-HC or 27-HC, but not with cholesterol, compared to vehicle control (Figure S7), suggesting that these oxysterols directly increase intracellular aSyn levels.

To finally confirm that the different cholesterol/metabolites are indeed taken up by neurons and to test whether their interaction with aSyn influences cholesterol metabolism, we performed a sterolomic analysis. Primary hippocampal neurons were treated with 10 µM 24S-HC or 27-HC, in the presence or absence of 500 nM extracellular aSyn, a concentration within the physiological range^43^. Sterolomic analysis revealed elevated intracellular levels of 24S-HC and 27-HC compared to vehicle-treated controls, indicating efficient cellular uptake of these oxysterols (Figure S8a and d). This was further supported by the increased abundance of their downstream metabolites (Figure S8b, c, e and f), suggesting active intracellular metabolism. Notably, the presence of extracellular aSyn led to an increase in the downstream metabolite of 24-HC, namely 7α,24-dihydroxycholest-4-en-3-one (7α,24-diHCO), but a modest reduction in 24S-HC itself and another downstream metabolite 7α,24-dihydroxycholesterol (7α,24-diHC) (Figure S8a, b and c). In contrast, extracellular aSyn enhanced the accumulation of 27-HC and its downstream metabolites, 7α,27-dihydroxycholesterol (7α,27-diHC) and 7α,27-dihydroxycholest-4-en-3-one (7α,27-diHCO)^44^ (Figure S8d, e, and f), suggesting that aSyn may facilitate the cellular uptake and metabolic processing of both 24S-HC and 27-HC, though with different magnitudes. Together, these findings support a direct molecular link between dysregulated cholesterol metabolism, aSyn pathology, and the progression of PD.

## Discussion

Neurodegenerative diseases are characterised by a progressive disruption of normal neuronal function, beginning with impaired synaptic transmission^45^ and culminating in a loss of synchronous neuronal firing. Given this trajectory, calcium signalling serves as a powerful readout for assessing the effects of cholesterol and its metabolites on neuronal function. Calcium dynamics are essential for synaptic plasticity and neurotransmitter release^24–26^, and are precisely regulated by a balance of cytosolic influx, efflux, and exchange with internal stores such as the ER and mitochondria^27^. Perturbations in this tightly controlled system can disrupt synaptic integrity and are increasingly recognised as contributing factors in the pathogenesis of neurodegenerative diseases^46^. Our study shows that 24S-HC and 27-HC dysregulate calcium dynamics in i^3^ neurons. Calcium imaging heatmaps revealed a broadening of calcium spikes following one day of treatment with 24S-HC and 27-HC, indicative of prolonged intracellular calcium elevations. Interestingly, increased dendritic calcium transient duration during activity behaviour has also been observed in APP/PS1 mice^47^. Additionally, both metabolites increased resting cytosolic calcium levels, consistent with previous findings in SK-N-BE and SH-SY5Y cells treated with these oxysterols^48,49^. Elevated cytosolic calcium level is a hallmark of several neurodegenerative diseases, including PD and AD^50^. For example, mutant aSyn and Aβ oligomers elevate cytosolic calcium by increasing plasma membrane ion permeability^51,52^. In our experiments, 24S-HC and 27-HC also decreased relative calcium spike amplitude, suggesting impaired calcium fluctuations, and inhibition or overactivation of calcium channels or transporters such as voltage-gated calcium channels and ryanodine receptors.

Following oxysterol treatment, we also observed elevated ER calcium levels in neuronal somas, paralleling reports of ER calcium overload in GBA1-linked PD fibroblasts^53^ and in familial AD due to presenilin mutations^54^. Since mitochondria sequester calcium from the ER at mitochondria-associated membranes (MAMs), ER calcium overload may propagate mitochondrial calcium stress, potentially inducing apoptosis or autophagy^55^. Additionally, 24S-HC and 27-HC altered calcium spike rate, although without a clear pattern, consistent with biphasic calcium spike rate changes seen in PD models^56^.

Our data further reveal an impaired calcium spike synchronisation, a feature critical for spike timing-dependent plasticity^57^ and particularly vital in the substantia nigra’s dopaminergic neurons, which exhibit regular calcium spikes and are highly sensitive to calcium overload^58^. Our findings thus suggest that 24S-HC and 27-HC significantly compromise neuronal network function by impairing both intracellular and intercellular calcium signalling.

Calcium homeostasis is tightly regulated by multiple intracellular organelles, including mitochondria, lysosomes, and the ER, all of which are known to be disrupted in neurodegenerative diseases. As such, organelle dysfunction may play a direct role in calcium dysregulation. To identify which organelle pathways could underlie the calcium signalling defects observed in our system, we treated i^3^ neurons with bafilomycin A1, MG-132, and FCCP, targeting lysosomes, the proteasome, and mitochondria, respectively. Each compound induced measurable alterations in calcium dynamics, underscoring the critical role of organelle function in maintaining calcium homeostasis. These results prompted a more detailed investigation into the specific contribution of each organelle to the observed calcium signalling defects.

We found that 24S-HC and 27-HC induce mitochondrial fragmentation in axons and dendrites, visualised through SIM. Mitochondrial dynamics, regulated by fission (Drp1) and fusion proteins (Mfn1, Mfn2, OPA1), are critical for neuronal energy demands^59,60^. Our findings suggest a shift toward fission, potentially disrupting synaptic density^61^ and transmission^62^. Similar mechanisms have been implicated in PD and AD through PINK1/LRRK2 mutations^63^ and altered Drp1 activity^64^.

Mitochondrial fragmentation also correlates with reduced membrane potential (ΔΨm) and increased reactive oxygen species (ROS), both seen in neurodegeneration^29^. Previous studies report that 24S-HC and 27-HC reduce ΔΨm^65–67^ and increase ROS^67–70^. We confirmed reduced spare respiratory capacity using the Seahorse Mito Stress Test, indicating mitochondrial dysfunction^71,72^—a severe threat to ATP-dependent neuronal processes, as highlighted by the defect to calcium signalling in i^3^ neurons upon oxysterol treatment.

Lysosomal dysfunction was evident following 27-HC treatment, which induced lysosomal enlargement and LMP, consistent with cathepsins leakage observed in murine myeloid immune cells and co-cultured SH-SY5Y cells and C6 cells^67,73^. These structural changes may reduce lysosomal mobility^74^ and function, impairing degradation and organelle clearance. Using the DQ-BSA assay, we found that both oxysterols reduced lysosomal degradation, which could impair mitophagy and exacerbate mitochondrial defects. These findings parallel disruptions in PINK1/Parkin-dependent mitophagy observed in PD^75,76^.

Lysosomes also regulate calcium storage and release^77^. LMP and reduced lysosomal acidification may cause unregulated calcium release^78^, potentially contributing to the calcium dysregulation observed. This was further supported by the observation that bafilomycin A1 induced calcium defects similar to those caused by oxysterol treatment^79^. Similarly, in GBA1 mutant fibroblasts and NPC1 disease, lysosomal calcium storage is impaired^53,80^, linking degradation deficits to calcium imbalance.

We also observed axonal swelling and disrupted ER–mitochondria interactions. Axonal swelling colocalised with enlarged mitochondria, suggesting compression of the tubular ER within axons. As the axonal ER is key for calcium buffering and neurotransmitter release^81,82^, such disruptions may promote synaptic and axonal degeneration^31,83^. Mitochondrial swelling may also underlie impaired MAM and calcium dysregulation. Prior studies link aSyn and DJ-1 mutations in PD to disrupted MAM integrity^84,85^, and 24S-HC esters have been shown to disrupt ER membranes in SH-SY5Y cells^86^. Future experiments measuring the ubiquitin-proteasome system activity and analysing MAMs using MAM-specific markers will be essential for validating these effects.

It is challenging to identify a single organelle as the primary cause of the calcium signalling defects we have observed. However, the elevated calcium concentrations seen across all oxysterol treatments are most closely replicated by bafilomycin A1 treatment. This is particularly noteworthy given that PD has also been characterised as a lysosomal storage disorder^87^. Lysosomes appear to play a central role in regulating the function and fate of multiple organelles, including mitochondria and the ER^88–90^, especially the tubular ER, which is the predominant ER form in axons. Our recent findings demonstrate that the tubular ER network is shaped and extended through the movement of lysosomes trafficked along microtubules^41^. Therefore, lysosomal enlargement and impaired trafficking to the periphery of the cells may disrupt the organisation of the tubular ER in axons, which in turn could impact mitochondrial fission and fusion^91^. Such disturbances may compromise the delivery of essential trophic factors, such as proteins, calcium, and lipids, to synapses^92^. Further experiments will be necessary to investigate these mechanisms in greater detail and to clarify the interdependence of these organelles in the context of oxysterol-induced dysfunction.

As an initial step toward exploring the connection between oxysterols and PD, we examined the molecular interactions between 24S-hydroxycholesterol (24S-HC), 27-hydroxycholesterol (27-HC), and alpha-synuclein (aSyn).This builds on previous studies suggesting that oxysterols can promote aSyn accumulation and aggregation, as well as earlier NMR findings indicating direct interactions between aSyn and cholesterol^33^. Our NMR analysis revealed that both 24S-HC and 27-HC bind specifically to the N-terminal region of aSyn with greater affinity than cholesterol. This higher binding affinity may enhance local aSyn concentration and promote conditions conducive to aggregation, thereby providing a potential molecular mechanism linking oxysterol accumulation to aSyn pathology in PD. This is further supported by experiments in SH-SY5Y cells overexpressing aSyn-YFP, where treatment with either oxysterol elevated aSyn-YFP levels. Whether this effect stems from direct molecular binding or results from transcriptional or translational upregulation remains to be clarified.

Additional evidence supporting a direct interaction comes from sterolomic analysis in primary neurons, where extracellular aSyn increased intracellular levels of 27-HC and its metabolites, 7α,27-diHC and 7α,27-diHCO. Interestingly, this aSyn-driven accumulation of 7α,27-diHC in neurons may correspond to the elevated levels of the same metabolite observed in the CSF of PD patients^19^. An increase in the terminal metabolite 7α,24-diHCO was also observed, accompanied by a modest reduction in 24S-HC and 7α,24-diHC. These findings point to a potential feedforward loop in which elevated 24S-HC and 27-HC levels promote aSyn uptake and aggregation, while accumulated aSyn enhances oxysterol internalisation, amplifying pathological interactions relevant to PD.

Despite these findings, several critical questions remain unresolved. Do 24S-HC and 27-HC preferentially bind to specific aSyn conformers or influence distinct aggregation pathways? Could these oxysterols alter aSyn’s membrane-or vesicle-binding properties in a manner that promotes its pathological accumulation? Additionally, might they compromise membrane integrity or interfere with intracellular trafficking, thereby facilitating increased uptake or mislocalisation of aSyn? It also remains unclear whether similar mechanisms apply to other amyloidogenic proteins, such as Aβ. Further studies are needed to elucidate these pathways and determine whether oxysterol–protein interactions play a direct and causative role in the pathogenesis of PD and other neurodegenerative disorders.

### Conclusion

In conclusion, our findings uncover a previously underappreciated role for cholesterol metabolites in initiating and perpetuating a cascade of cellular dysfunction that spans calcium signalling, mitochondrial integrity, lysosomal capacity, and ER homeostasis. The convergence of these disruptions suggests that oxysterols act as central mediators of organelle crosstalk failure, amplifying neuronal stress through a self-reinforcing cycle of dysfunction. Coupled with their capacity to promote protein aggregation, 24S-HC and 27-HC emerge as potent contributors to neurodegenerative pathology. These insights position cholesterol metabolites not merely as bystanders, but as active drivers of disease progression. Targeting their accumulation or downstream effects may therefore offer a novel and urgently needed therapeutic avenue for halting or reversing neurodegeneration.

## Acknowledgements

G.S.K.S. acknowledges funding from the Wellcome Trust (065807/Z/01/Z and 203249/Z/16/Z), the UK Medical Re-search Council (MRC) (MR/K02292X/1), Alzheimer Research UK (ARUK-PG013-14), Michael J Fox Foundation (16238 and 022159), and Infinitus China Ltd. C.F.K. acknowledges funding from the UK Engineering and Physical Sciences Re-search Council (EPSRC) (EP/L015889/1 and EP/H018301/1), the Wellcome Trust (3-3249/Z/16/Z and 089703/Z/09/Z), the MRC (MR/K015850/1), and Infinitus China Ltd. Y.W. and W.G. acknowledge funding from UKRI MR/X012387/1 and BB/S019588/1.

