## Supplementary Information for "Brain cholesterol metabolites cause significant neurodegeneration in human iPSC-derived neurons"

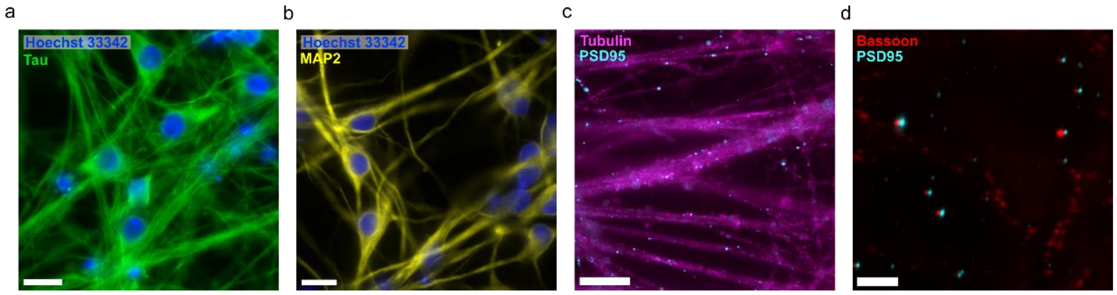


**Figure S1. Differentiated i$^3$ neurons display a neuronal phenotype after 25 days in vitro.** i$^3$ neurons were fixed at 25 DIV and stained with Hoechst 33342 (a nucleic acid stain), Tau, microtubule associated protein 2 (MAP2, a neuronal dendritic marker), beta-tubulin, postsynaptic density 95 (PSD95, a postsynaptic marker), and Bassoon (a presynaptic marker). Scale bars: 20 µm for **(a) - (c)**, and 5 µm for **(d)**.

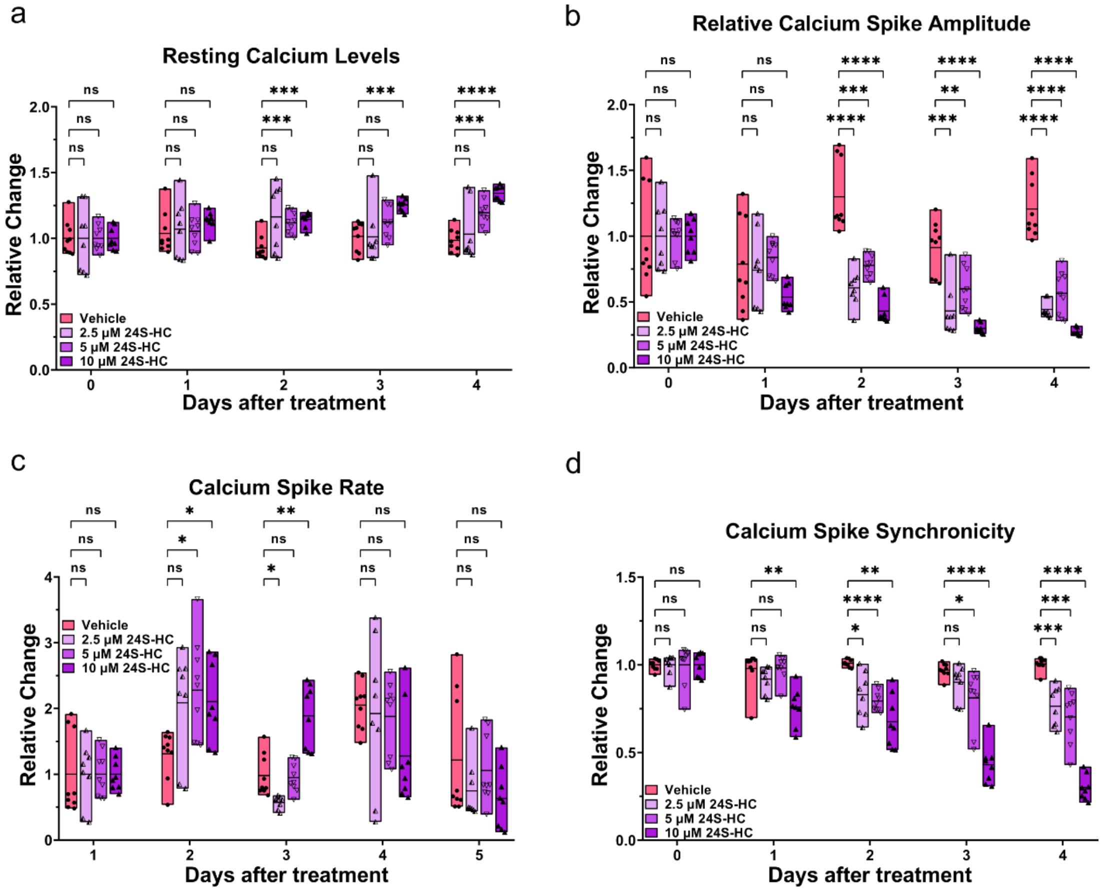


**Figure S2. 24S-HC impairs calcium signalling in i$^3$ neurons in a dose-dependent manner. (a) - (d)** The relative change in resting calcium levels ($F_0$), relative calcium spike amplitude (peak $\Delta F/F_0$), calcium spike rate and calcium spike synchronicity of i$^3$ neurons treated with 2.5, 5 or 10 µM 24S-HC over 4 days. Data were normalised to

Day 0 (pretreatment) and presented as floating bar plots (min to max), with a line indicating the mean. One field of view was imaged per well, and each data point represents a single well. Two-way ANOVA with Dunnett's multiple comparisons tests, where ns is not significant, * is p<0.05 and **** is p<0.0001. Data from N = 3 independent experiments were quantified.

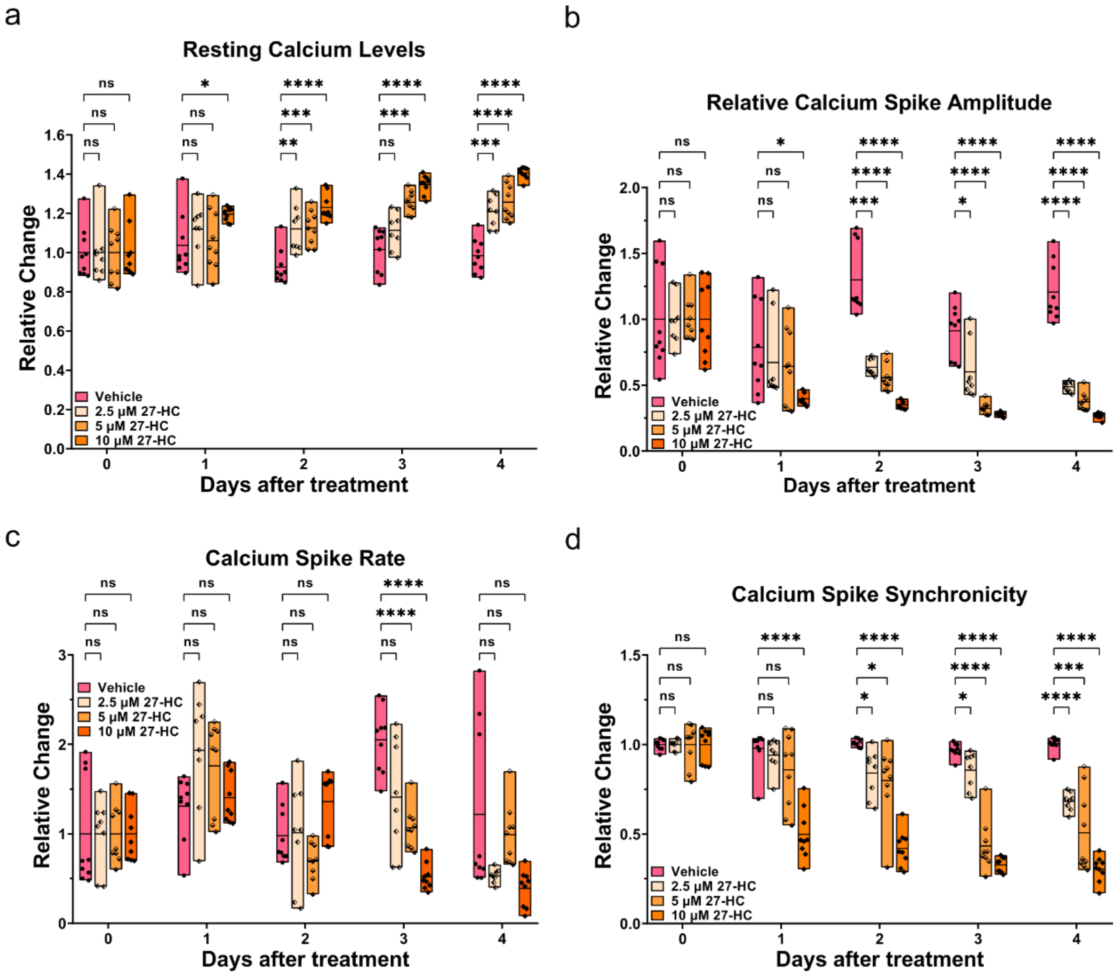


**Figure S3. 27-HC impairs calcium signalling in i[3] neurons in a dose-dependent manner. (a) - (d)** The relative change in resting calcium levels ($F_0$), relative calcium spike amplitude (peak $\Delta F/F_0$), calcium spike rate and calcium spike synchronicity of i[3] neurons treated with 2.5, 5 or 10 µM 27-HC over 4 days. Data were normalised to Day 0 (pretreatment) and presented as floating bar plots (min to max), with a line indicating the mean. One field of view was imaged per well, and each data point represents a single well. Two-way ANOVA with Dunnett's multiple comparisons tests, where ns is not significant, * is p<0.05 and **** is p<0.0001. Data from N = 3 independent experiments were quantified.

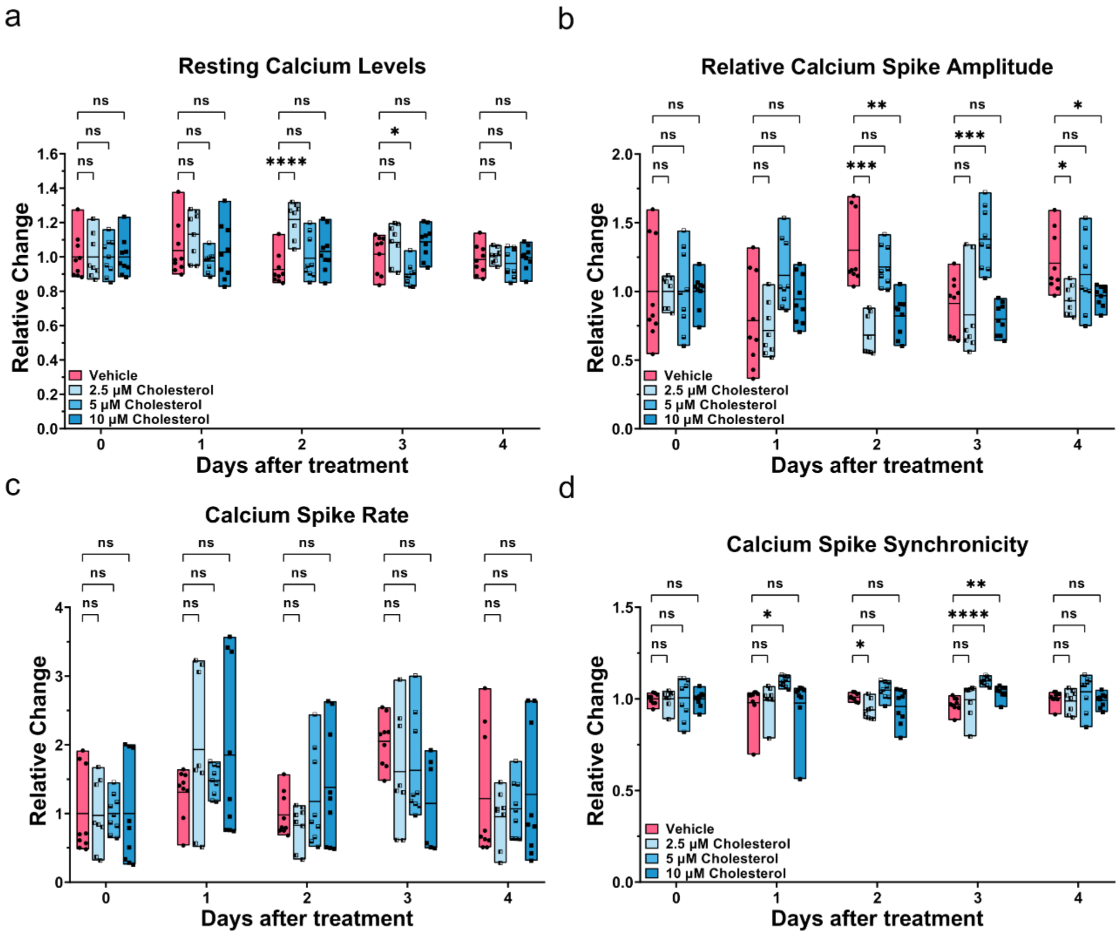


**Figure S4. Cholesterol does not exhibit a clear dose-dependent effect on calcium signalling in i[3] neurons. (a) - (d)** The relative change in resting calcium levels ($F_0$), relative calcium spike amplitude (peak $\Delta F/F_0$), calcium spike rate and calcium spike synchronicity of i[3] neurons treated with 2.5, 5 or 10 µM cholesterol over 4 days. Data were normalised to Day 0 (pretreatment) and presented as floating bar plots (min to max), with a line indicating the mean. One field of view was imaged per well, and each data point represents a single well. **(a)**, **(b)** and **(d)** were analysed with two-way ANOVA with Dunnett's multiple comparisons tests, while **(c)** was analysed with mixed-effects analysis with Dunnett's multiple comparisons tests. ns is not significant, * is $p<0.05$ and **** is $p<0.0001$. Data from N = 3 independent experiments were quantified.

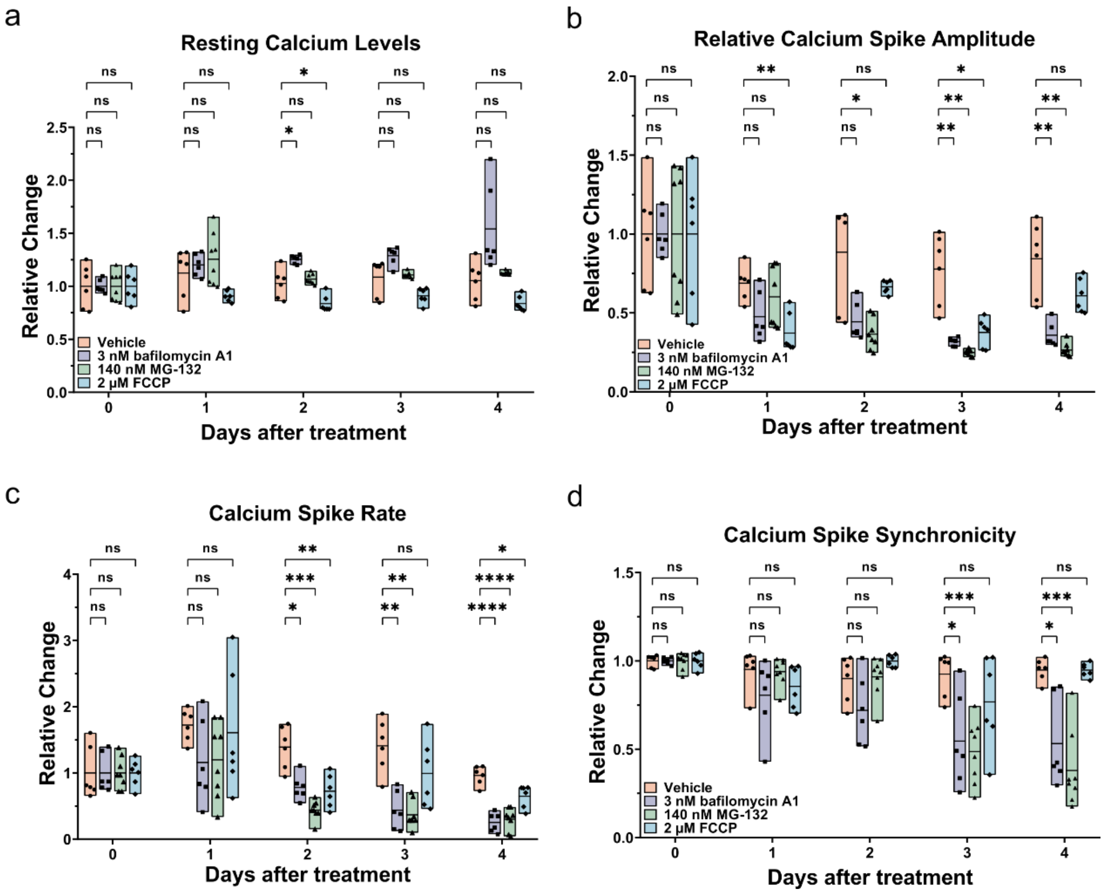


**Figure S5. Pharmacological inhibitors affect calcium signalling in i$^3$ neurons. (a) - (d)** The relative change in resting calcium levels ($F_0$), relative calcium spike amplitude (peak $\Delta F/F_0$), spike rate and calcium spike synchronicity of i$^3$ neurons treated with 3 nM bafilomycin A1, 140 nM MG-132 and 2 μM FCCP over 4 days. Data were normalised to Day 0 (pretreatment) and presented as floating bar plots (min to max), with a line indicating the mean. One field of view was imaged per well, and each data point represents a single well. Two-way ANOVA with Dunnett's multiple comparisons tests, where ns is not significant, * is $p<0.05$ and **** is $p<0.0001$. Data from $N = 3$ independent experiments were quantified.

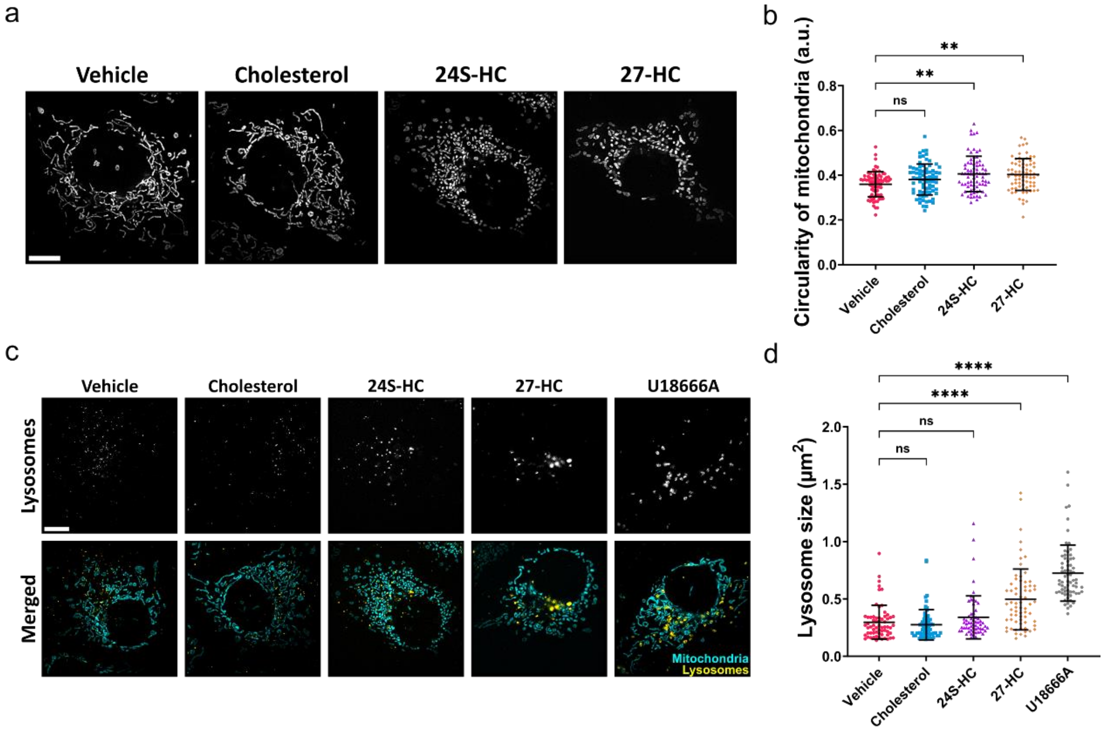


**Figure S6. 24S-HC and 27-HC significantly affect mitochondria and lysosome morphology in COS-7 cells. (a)** Representative SIM images of mitochondria in COS-7 cells treated with 10 µM cholesterol, 24S-HC or 27-HC for 1 day. Scale bar = 10 µm. **(b)** The circularity of mitochondria is presented as means ± S.D. Each data point represents an individual image. Kruskal-Wallis test with Dunn's multiple comparisons, where ns is not significant and ** is $p<0.01$. Data from N = 3 independent experiments were quantified. **(c)** Representative SIM images of lysosomes in COS-7 cells treated with 10 µM cholesterol, 24S-HC or 27-HC for 1 day. Scale bar = 10 µm. **(d)** The size of lysosomes is presented as means ± S.D. Each data point represents an individual image. Kruskal-Wallis test with Dunn's multiple comparisons, where ns is not significant and **** is $p<0.0001$. Data from N = 3 independent experiments were quantified.

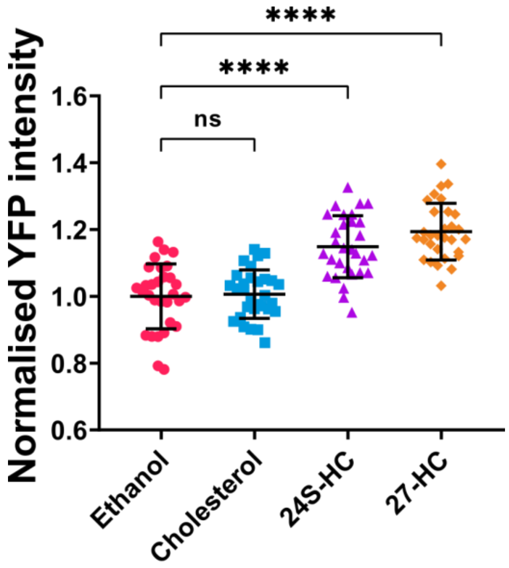


**Figure S7. 24S-HC and 27-HC increase aSyn levels in SH-SY5Y cells.** SH-SY5Y cells overexpressing aSyn-YFP were treated with 10 μM cholesterol, 24S-HC or 27-HC for 2 days. The normalised YFP intensity is presented as means ± S.D. Each data point represents an individual image. One-way ANOVA tests with Dunnett's multiple comparison, where ns is not significant and **** is p<0.0001. N = 3 independent experiments were quantified.

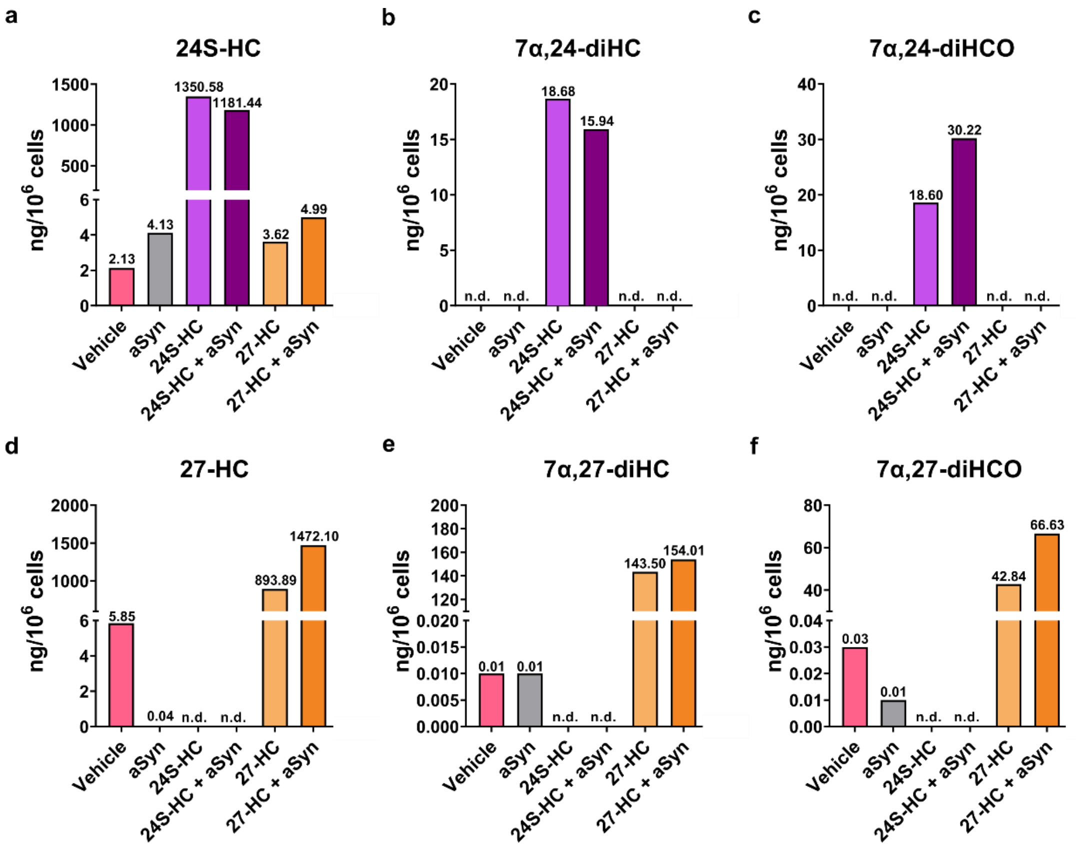


**Figure S8. 24S-HC and 27-HC are effectively taken up by neuronal cells.**

Sterolomic analysis of primary hippocampal neurons treated with 10 µM 24S-HC or 27-HC, in the presence or absence of 500 nM extracellular aSyn. The X-axis indicates the treatment conditions, and the graph titles specify the lipid analysed. Numerical values are displayed above each bar, and 'n.d.' indicates not detected. **(a), (b) and (c)** 24S-HC and its downstream metabolites, 7α,24-diHC and 7α,24-diHCO. **(d), (e) and (f)** 27-HC and its downstream metabolites, 7α,27-diHC and 7α,27- diHCO. Abbreviations: HC, hydroxycholesterol; diHC, dihydroxycholesterol; diHCO, dihydroxycholest-4-en-3-one.

### Supplementary methods

**Table S1. Reagents used in the supplementary.**

| Reagent Or Resource | Source | Identifier | Dilution |
|---|---|---|---|
| **Oxysterol analysis** | | | |
| [25,25,25,26,26,26-$^2H_6$]24R/S-Hydroxycholesterol | Avanti Polar Lipids (now Avanti Research) | LM4110 | - |
| 27-Hydroxycholesterol ((25R)26-Hydroxycholesterol) | Avanti Polar Lipids (now Avanti Research) | 700021P | - |
| 7α,24S-Dihydroxycholesterol | Avanti Polar Lipids (now Avanti Research) | 700096P | - |
| [25,25,25,26,26,26-[$^2H_6$]7α,25-Dihydroxycholesterol | Avanti Polar Lipids (now Avanti Research) | 111117P | - |
| 7α,27-Dihydroxycholesterol (7α,(25R)26-Dihydroxycholesterol) | Avanti Polar Lipids (now Avanti Research) | 700024P | - |
| [24,24,27,27,27-$^2H_5$]3β-Hydroxycholest-5-en-(25R)26-oic acid | Avanti Polar Lipids (now Avanti Research) | 700151P | - |
| Cholesterol oxidase from Streptomyces sp | Merck | C8649 | - |
| Girard P reagent | Tokyo Chemical Industry | G0030 | - |

| | | | |
|---|---|---|---|
| **SH-SY5Y cell culture** | | | |
| Minimum Essential Medium Eagle (MEM) | Sigma-Aldrich | M2279-100ML | - |
| Nutrient Mixture F-12 Ham | Sigma-Aldrich | N4888-500ML | - |
| **Dyes** | | | |
| Dextran, Alexa Fluor™ 594; 10,000 MW, Anionic | Invitrogen | D22913 | - |
| **Antibodies** | | | |
| MAP2a + MAP2b (MT-07) Antibody | Abcam | ab36447 | 1:100 |
| Tau Monoclonal Antibody (T46) | Thermo Fisher | 13-6400 | 1:200 |
| Anti-beta Tubulin antibody | Abcam | ab15568 | 1:200 |
| Bassoon (D63B6) | Cell signalling | 6897 | 1:400 |
| Anti-PSD95 antibody | Abcam | ab99009 | 1:200 |
| Hoechst 33342 | Invitrogen | H3570 | 1:1000 |
| Goat anti-Mouse IgG (H+L) Highly Cross-Adsorbed Secondary Antibody, Alexa Fluor™ 647 (MAP2) | Invitrogen | A-21236 | 1:100 |
| Goat anti-Mouse IgG (H+L) Highly Cross-Adsorbed Secondary Antibody, Alexa Fluor™ 568 (Tau, PSD95) | Invitrogen | A-11031 | 1:100 |
| Goat anti-Rabbit IgG (H+L) Highly Cross-Adsorbed Secondary Antibody, Alexa Fluor™ 647 (Tubulin) | Invitrogen | A-21245 | 1:100 |
| Goat anti-Rabbit IgG (H+L) Cross-Adsorbed Secondary Antibody, Alexa Fluor™ 647 (Bassoon) | Invitrogen | A-21244 | 1:100 |

### SH-SY5Y cell culture

SH-SY5Y cells expressing aSyn-yellow fluorescent protein (YFP) were engineered by lentiviral transfection as previously described[1]. Cells were cultured at 37°C with a 5% $CO_2$ atmosphere. The cell culture media is composed of 41.5 v/v% minimal essential medium (MEM), 41.5 v/v% nutrient mixture Ham's F-12, 15 v/v% fetal bovine serum

(FBS), 2 mM GlutaMAX, and 1X antibiotic-antimycotic. SH-SY5Y cells were passaged when they reached 80-90% confluence. For widefield microscopy imaging, SH-SY5Y cells were plated in µ-slide 8 well high glass-bottom plates (ibidi GmbH) with about 20,000 cells per well, and treated with different concentrations of cholesterol, 24S-HC or 27-HC for 2 days.

### Primary hippocampal neuronal culture and treatment

Primary hippocampal neurons were isolated from postnatal day 2 (P2) Sprague-Dawley rats (Charles River) as previously described[2]. 1 million cells were plated in each 35 mm dish (MatTek) coated with 0.01% poly-l-lysine. The cells were cultured for 21 days in 2 mL Neurobasal medium supplemented with 1X B-27 supplement, 0.5 mM GlutaMAX supplement and 1X antibiotic-antimycotic. Half of the media was changed once a week. At 21 DIV, cells were treated with 10 µM 24S-HC or 27-HC, in the presence or absence of 500 nM extracellular aSyn, which was expressed and purified as described previously[3], and incubated for 3 days. After treatment, cells were washed twice with cold PBS, collected with cold PBS and centrifuged at 13,000 rpm for 15 seconds. Supernatant was removed and the cell pellet was frozen in dry ice for the following lipidomic measurement.

### $i^3$ neuron immunolabeling

$i^3$ neurons were fixed with 4% formaldehyde solution in PBS at 26 DIV. After being washed with PBS 3 times, cells were permeabilized with 0.05% PBS-Tween20 (PBST) for 5 minutes and then blocked with 5% donkey serum in PBST for 1 h. Primary antibodies were diluted in PBST and incubated with the cells for 1 h at room temperature (Table S1), followed by washing with PBST 3 times. Secondary antibodies were diluted in PBST and incubated with the cells for 1 h at room temperature (Table S1), followed by washing with PBST 3 times. To label DNA, Hoechst 33342 was added for 40 minutes and washed once with PBS.

### Lysosome staining in COS-7 cells

Lysosome labeling was performed using the pulse-chase method. On the day before imaging, 0.1 mg/mL Dextran, Alexa Fluor™ 594; 10,000 MW, Anionic was added to the cells for 4 h, followed by washing with the culture media once and incubating with treatment for 20 h.

### Widefield microscopy

Widefield imaging was performed with an automated custom-built widefield microscope. 4-wavelength high-power LED source (LED4D067, Thorlabs), stage

(Prior), frame (IX83, Olympus), Z drift compensator (IX3-ZDC2, Olympus) and an sCMOS camera (Zyla sCMOS, Andor) were controlled by Micro-Manager. Filter cubes for Hoechst 33342 (filter set 49000-ET-DAPI, Chroma), YFP (filter set 49003-ET-EYFP, Chroma, emission filter replaced by 600LP, Semrock), Alexa Fluor 568 (filter set 49008-ET-mCherry, Texas Red, Chroma) and Alexa Fluor 647 (excitation filter 628/40, dichroic beamsplitter Di02-R635, emission filter 708/75, Semrock) were used. For aSyn-YFP and immunolabeling, images were captured with a 20×/0.45 NA air (LUCPlanFL N, Olympus), or a 60×/1.42 NA oil (PlanApo U, Olympus) objective lens, respectively.

### Oxysterol analysis

The method for sterol extraction, derivatization, and LC-MS has been previously described in detail[4]. Briefly, oxysterols in cell pellets were extracted in ethanol containing a cocktail of isotope-labelled standards including [26,26,26,27,27,27-$^{2}H_{6}$]24R/S-hydroxycholesterol ([$^{2}H_{6}$]24R/S-HC), [25,25,25,26,26,26-$^{2}H_{6}$]7α,25-dihydroxycholesterol ([$^{2}H_{6}$]7α,25-diHC) and [24,24,27,27,27-$^{2}H_{5}$]3β-hydroxycholest-5-en-(25R)26-oic acid ([$^{2}H_{5}$]3β-HCA). Oxysterols were separated from cholesterol by $C_{18}$ solid-phase extraction and derivatized (i) with [$^{2}H_{5}$] Girard P reagent following cholesterol oxidase treatment to oxidase 3β-hydroxy to 3-oxo groups or (ii) with [$^{2}H_{0}$] Girard P in the absence of cholesterol oxidase, in this case derivatising oxysterols with a natural 3-oxo group. The derivatised oxysterols were analysed using liquid chromatography-mass spectrometry with multistage fragmentation (LC-$MS^{n}$) on an Orbitrap IQX mass spectrometer coupled with Ultimate 3000 LC system (ThermoFisher Scientific). The identifications were based on exact mass, retention time and by comparing $MS^{3}$ fragmentation spectra to authentic standards. Oxysterols were quantified by the isotope dilution method.
